# Inducible nitric oxide synthase (iNOS) is necessary for GBP-mediated *T. gondii* restriction in murine macrophages via vacuole nitration and intravacuolar network collapse

**DOI:** 10.1101/2023.07.24.549965

**Authors:** Xiao-Yu Zhao, Samantha L. Lempke, Jan C. Urbán Arroyo, Bocheng Yin, Nadia K. Holness, Jamison Smiley, Sarah E. Ewald

## Abstract

*Toxoplasma gondii* is an obligate intracellular, protozoan pathogen of rodents and humans. *T. gondii’s* ability to grow within cells and evade cell-autonomous immunity depends on the integrity of the parasitophorous vacuole (PV). Interferon-inducible guanylate binding proteins (GBPs) are central mediators of *T. gondii* clearance, however, the precise mechanism linking GBP recruitment to the PV and *T. gondii* restriction is not clear. This knowledge gap is linked to heterogenous GBP-targeting across a population of vacuoles and the lack of tools to selectively purify the intact PV. To identify mediators of parasite clearance associated with GBP2-positive vacuoles, we employed a novel protein discovery tool automated spatially targeted optical micro proteomics (autoSTOMP). This approach identified inducible nitric oxide synthetase (iNOS) enriched at levels similar to the GBPs in infected bone marrow-derived myeloid cells. iNOS expression on myeloid cells was necessary for mice to control *T. gondii* growth in vivo and survive acute infection. *T. gondii* infection of IFNγ-primed macrophage was sufficient to robustly induce iNOS expression. iNOS restricted *T. gondii* infection through nitric oxide synthesis rather than arginine depletion, leading to robust and selective nitration of the PV. Optimal parasite restriction by iNOS and vacuole nitration depended on the chromosome 3 GBPs. Notably, GBP2 recruitment and ruffling of the PV membrane occurred in iNOS knockouts, however, these vacuoles contained dividing parasites. iNOS activity was necessary for the collapse of the intravacuolar network of nanotubular membranes which connects parasites to each other and the host cytosol. Based on these data we conclude reactive nitrogen species generated by iNOS cooperate with the chromosome 3 GBPs to target distinct biology of the PV that are necessary for optimal parasite clearance in murine myeloid cells.

## Introduction

*Toxoplasma gondii* is an obligate intracellular parasite with a remarkably broad intermediate host range that includes humans and mice. Although *T. gondii* infects approximately one-third of the human population, the infection is often asymptomatic^1^. Severe complications emerge in immuno-compromised individuals or infection of the fetus, underscoring the importance of a functional immune response to control *T. gondii* infection^2^.

The parasite’s ability to infect and grow within most nucleated cell types in hundreds of host species is related to the formation and maintenance of the parasitophorous vacuole (PV). The PV membrane is formed from the host plasma membrane during *T. gondii* invasion and connected to the parasite by an intra-vacuolar network (IVN) of host-derived lipid nano-tubes^3^. The parasite uses secreted effectors to maintain the PV and the IVN membranes, recruit nutrients to the vacuole, and evade immune recognition^4^. Formation of the PV facilitates the evasion of Toll-like receptors TLR7, -8, -9^5, 6^ and -11 (TLR11 is a pseudogene in humans), which recognize phagocytosed, damaged parasites^7^. Disruption of the gut epithelium during oral infection leads to TLR2, -4, and -9 signaling in response to commensal microbiota^5, 8, 9^. TLR signaling through NF- κB produces interleukin 12 (IL-12), which is critical for IFNγ production^7, 10, 11^. IFNγ is necessary for both the protective CD8 T cell response and for cell-autonomous control of parasite growth in human and murine cells^12^.

IFNγ receptor signaling through STAT1 transcriptionally upregulates interferon-inducible GTPases (IIGs) which survey cells for damaged or non-self-membranes and target them for removal^13–15^. In mice, the p47 immunity-related GTPases (IRG)a6, -b6 and -b10 are recruited to the PV membrane^16, 17^, under the regulatory control of IRGM1, -2 and -3^13, 14^. In addition, the p65 small guanylate-binding proteins (GBPs) encoded on chromosome 3 (GBP1-3, 5, and 7)^18^ and chromosome 5 (GBPs 6, 9)^19, 20^ are recruited to PV membrane and play partially redundant roles in vacuole disruption and parasite clearance. Mice deficient in regulatory *Irgm1*^21^, *Irgm2*^22, 23^, or *Irgm3*^24^ or chromosome 3 GBPs^18^ succumb to acute parasite overgrowth, which is partially recapitulated with delayed kinetics by *Irgd*^21^, *Irgb6*^25^*, Irga6*^26^, *Gbp1*^27^, *Gbp2*^28^ or *Gbp7*^29^ deficiency. This is also reflected in tissue culture where deleting the IRGs or chromosome 3 GBPs results in a partial rescue of parasite growth, depending on cell type^18, 25–27^. The IIGs have high within-host allelic diversity, consistent with positive selection imparted by pathogen infection, as high variation across species^30–32^. In human cells, the functional role of IRGs is not conserved; and GBP1, -2 and -5 have been shown to be necessary for IFNγ-mediated *T. gondii* clearance, although other human GBPs are also recruited to the PV membrane^33, 34^.

Despite advances in our understanding of the IIG family members and their mechanism of recruitment to the vacuole, a complete mechanism of *T. gondii* vacuole disruption and killing is lacking. Live imaging studies have shown that IRGb6 recruitment to the PV is followed by the re-localization of fluorescent proteins from the host cytosol to the parasite then parasite rounding (a measure of death)^16^. Mouse IRGb6^25^, GBP1^27^ and chromosome 3 GBPs^18^ are associated with PV membrane ‘ruffling’ and discontinuity by transmission electron microscopy. Human GBP1 (the homolog of mouse GBP2) positive vacuoles are associated with permeability to cytosolic dye and leakage of parasite nucleic acids to the cytosol compared to GBP1-negative vacuoles^33^. These studies have been taken as evidence that GBP recruitment and membrane deformation lead to parasite clearance via mechanisms analogous to what has been shown for bacterial outer membrane disruption^35, 36^. However, *T. gondii* resides in a vacuole with a fundamentally distinct origin from intracellular bacterial pathogens (PV versus modified phagosome) and uses cell membrane biology that is more similar to the host than bacteria, indicating that there are holes in our understanding of the mechanism of parasite clearance.

Heterogeneous IIG targeting to the PV has been a major challenge to understanding the mechanism of cell-autonomous *T. gondii* clearance. The frequency of IIG-positive vacuoles ranges from 30-60% of all PV depending on the identity of the IIG, host and parasite cell type^18, 19, 27, 28, 33^. Tools to isolate the PV are limited. Subcellular fractionation does not preserve the intact PV membrane^37^. A proximity biotinylation tool to study the PV proteome has been developed, however, it cannot differentiate between IIG-targeted and IIG-negative vacuoles in the population^38^. An additional hurdle is that IRG and GBP activity is also spatially regulated. A single host cell contains both inactive pools of the GTPases localized to vesicles, Golgi, or ER and active GTPase pools targeted to pathogen-associated membranes or damaged-self membranes^14, 15^.

Here we leverage the heterogeneity in IIG recruitment to the PV for comparative proteomics using **auto**mated **s**patially targeted **o**ptical **m**icro-**p**roteomics (autoSTOMP)^39, 40^. This led to the discovery that iNOS expression in murine macrophages and dendritic cells is necessary for efficient, parasite restriction by GBPs recruited to the PV membrane. Rather than mediating parasite clearance by arginine deprivation, iNOS activation leads to nitration of the vacuole by reactive nitrogen species, causing the collapse of the inner vacuolar network (IVN) which is necessary for nutrient import from the host and transport of parasite effectors to the host cytosol^3, 41^. The efficiency of IVN collapse and iNOS-mediated killing is dependent on the chromosome 3 GBPs, indicating that these two innate immune effector pathways collaborate to mediate vacuole-autonomous parasite clearance.

## Results

### IFNγ treatment leads to heterogeneous GBP2 recruitment and efficient *T. gondii* restriction in BM-MoDCs

To assess the role of immunological stress on *T. gondii* restriction, BM-MoDCs were incubated with media alone, IFNγ and/or the TLR2 ligand Pam3CSK4. BM-MoDCs were then infected with the type II *T. gondii* strain Me49 which is sensitive to IIG-mediated restriction and engineered to express both GFP and luciferase. *T. gondii* luciferase intensity increased over a 14-hour infection in unstimulated and TLR2-stimulated BM-MoDCs (Figure 1A, black and red). In contrast, BM-MoDCs treated with IFNγ alone or IFNγ and Pam3CSK4 efficiently restricted *T. gondii* growth by 14 hours post-infection (Figure 1A blue and violet). The IFNγ-dependent reduction in luciferase signal corresponded with decreased number of parasite vacuoles by 14 hours post-infection (Figure 1B).

**Figure 1.**
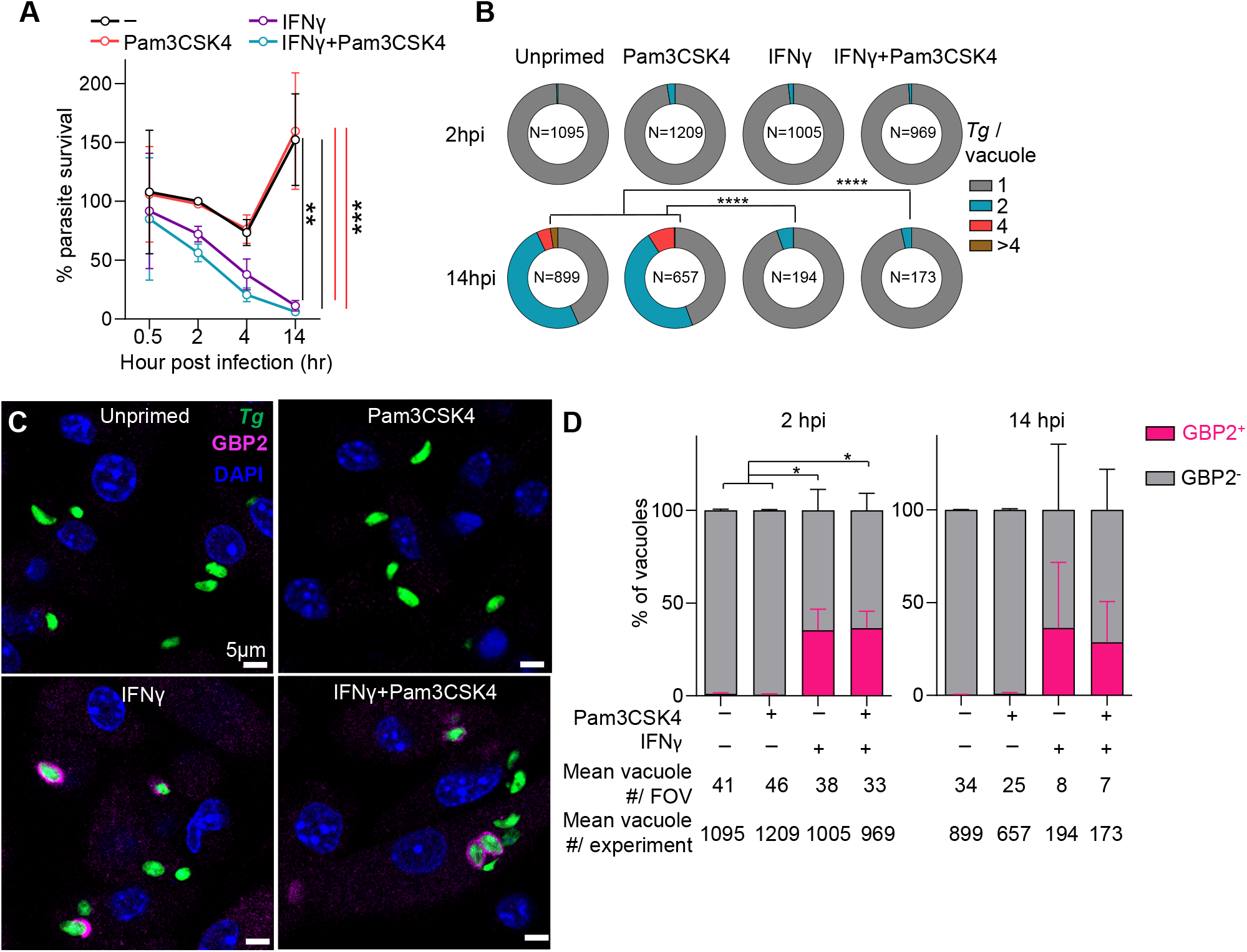
IFNγ stimulation leads to heterogenous GBP2 recruitment to the *T. gondii* vacuole in BM-MoDCs. A-D. B6 mouse bone marrow monocyte/dendritic cells (BM-MoDCs) differentiated with GM-CSF were pretreated for 20 hours with media, 70ng/ml IFNγ and/or 100ng/ml of Pam3CSK4 for the final 3 hours before infection. BM-MoDCs were infected with *T. gondii* expressing GFP and luciferase (Me49GFP-Luc *Tg*) at MOI 1.5. **A**, At 0.5, 2, 4, or 14-hours post-infection (hpi) parasite load was quantified by luciferase assay and plotted as relative light units (RLU) normalized to unstimulated, infected cells at 2 hpi. N=3 independent experiments. **B**, The number of *Tg* per vacuole (1, 2, 4 or greater than 4 parasites) was quantified by microscopy and plotted as the proportion of all vacuoles. **C,** GBP2 (magenta) localization to the *Tg* vacuole (green) was determined by immunofluorescence microscopy, counterstained with DAPI (blue). **D,** The number of GBP2 positive vacuoles (magenta) was quantified and plotted as the percent of all vacuoles. N=3 independent experiments. Scale bar, 5 μm. Error bars represent Mean±SEM by one-way ANOVA (**A**) or 2-way ANOVA (**B**, **D**) with Tukey post hoc test. *, p≤0.05; **, p≤0.01; ***, p≤0.001; ****, p≤0.0001.

GBP2 co-localized with 40% of *T. gondii* vacuoles by two hours post-infection in BM- MoDCs treated with IFNγ or IFNγ and Pam3CSK4 (Figure 1C), but not in unstimulated or Pam3CSK4-only stimulated conditions. Cytosolic puncta of GBP2 were also observed, consistent with the presence of a non-targeted, vesicular pool of GBP2^19, 20, 42^. While the total number of parasites decreased over time (Figure 1A-B) the proportion of GBP2^+^ and GBP2^-^ vacuoles was similar at 2- and 14-hours post-infection (Figure 1D). This heterogeneity in GBP targeting is consistent with previous observations in mouse macrophages^27^ and embryonic fibroblasts (MEFs)^28^.

### Inducible nitric oxide synthase (iNOS) is enriched at GBP2-targeted vacuoles in IFNγ- treated BM-MoDCs by automated spatially targeted optical micro proteomics (autoSTOMP)

IIG biology is regulated transcriptionally and post-translationally via localization, phosphorylation and prenylation^13, 14, 43–47^. We expected host effectors regulating vacuole disruption and parasite clearance to be differentially enriched on parasite vacuoles under restriction conditions compared to non-restrictive conditions, however, the presence of multiple pools of IIG (Figure 1C) in the cell^16, 18, 21, 26–28, 33, 42^ and heterogeneity in IIG targeting (Figure 1D) posed a barrier to identifying mediators of *T. gondii* targeting and clearance with bulk-analysis or proximity-mediated proteomic and transcriptomic tools^38, 48^. To identify regulators of IIG-mediated parasite clearance we used automated spatially targeted optical micro proteomics (autoSTOMP)^39, 40^ to image and selectively photo-biotinylate proteins in the ∼1μm region surrounding the *T. gondii* signal in unstimulated, Pam3CSK4, IFNγ or IFNγ and Pam3CSK4 stimulated BM-MoDCs (Figure 2A, B, PVM). On replicate coverslips, autoSTOMP was used to selectively biotinylate regions where GBP2 was adjacent to or colocalized with the *T. gondii* signal in the IFNγ or IFNγ and Pam3CSK4 stimulated conditions (Figure 2A, GBP2^+^). Biotinylated proteins were streptavidin precipitated and identified by liquid chromatography-mass spectrometry (LC-MS) with label-free quantification.

**Figure 2.**
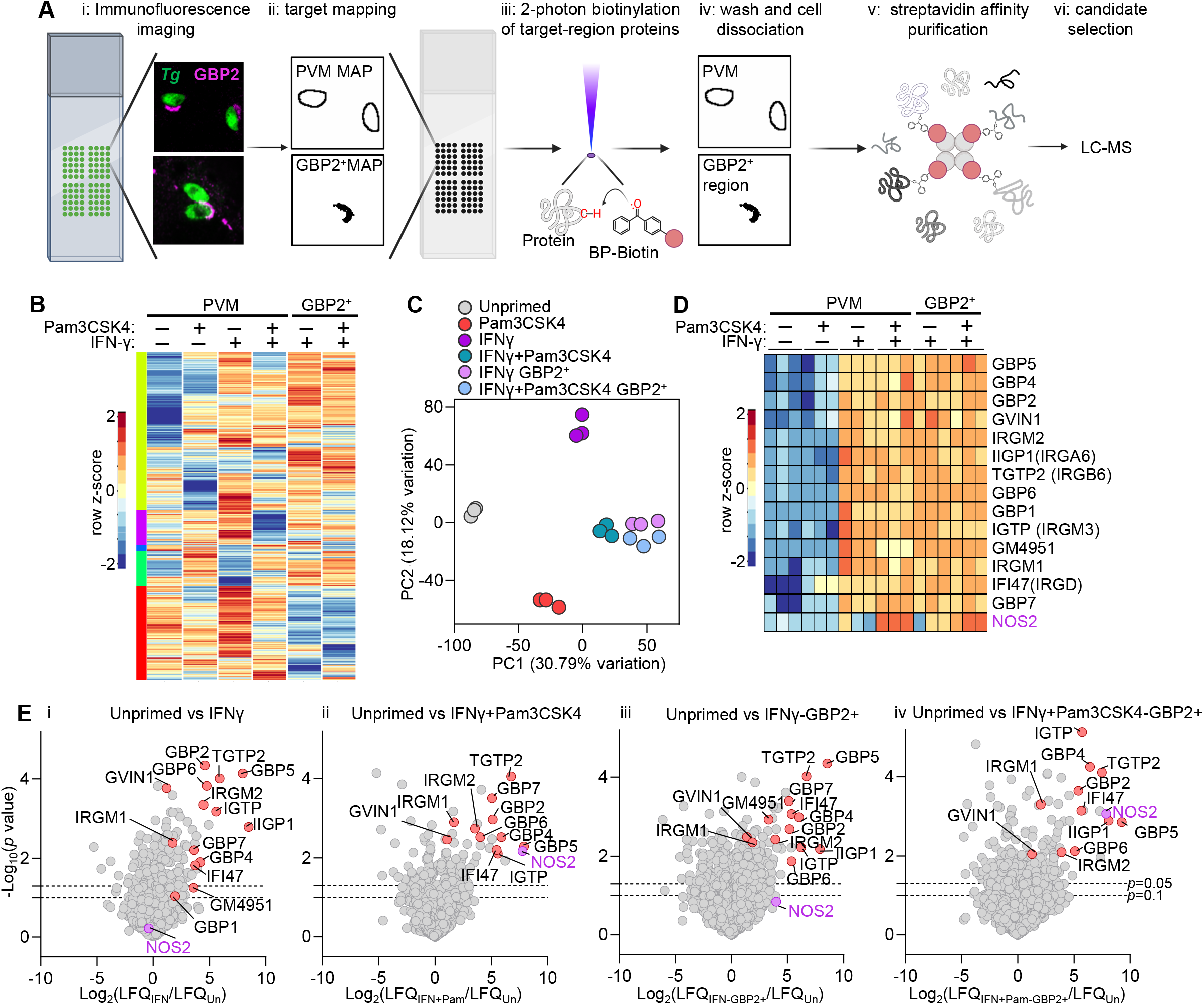
AutoSTOMP identifies inducible nitric oxide synthase (iNOS) enriched near parasite vacuoles in IFNγ and Pam3CSK4-treated BM-MoDCs. A. BM-MoDCs were prepared for autoSTOMP as described in Figure 1. i, At 2 hpi cells were fixed and stained for immunofluorescence imaging with *Tg* and GBP2-specific antibodies. ii, Images were used to identify the pixel coordinates of the regions containing the parasite vacuole membrane and immediate cytosol (PVM MAP) or regions where GBP2^+^ co-localized with the vacuole (GBP2^+^ MAP). iii, MAP files were used to guide 2-photon laser excitation to photo-crosslinking BP-biotin to any protein in the target regions. (iv) Unconjugated BP-biotin was washed away and cell lysates were prepared for (v) streptavidin bead affinity purification. vi, biotinylated proteins were identified by LC-MS and label-free quantification (LFQ). **B**. 2,070 mouse proteins were identified across 6 autoSTOMP conditions. Heatmap represents the row Z-score of each protein’s log_2_LFQ, averaged across N=3 biological replicates per condition, clustered by Z-score similarity across conditions (color blocks at left). **C,** PCA analysis of autoSTOMP replicates. **D,** IFN-inducible GTPases (IIGs) and iNOS identified in autoSTOMP, cropped heat map from (**B**). **E,** Pairwise comparison of proteins enriched in the unprimed PVM region (left, negative values) relative to IFNγ or IFNγ+Pam3CSK4 PVM regions or the subset of GBP2^+^ vacuoles. IIGs, red; iNOS, violet. Plots showing –log_10_(*p* value) from Student’s t test and log_2_LFQ differences between each comparison. Dotted lines indicated *p*=0.1 and *p*=0.05 respectively.

2,070 host proteins were identified across the six PVM and GBP2^+^ samples (Figure 2B, Supplementary Table S1). Principal component analysis confirmed that biological replicates were more similar to each other than differentially stimulated conditions collected and analyzed on the same days (Figure 2C). Of note, the GBP2^+^ ‘IFNγ’ and ‘IFNγ and Pam3CSK4’ stimulated conditions clustered tightly (Figure 2D pink, blue) indicating that IIG positive vacuoles have a conserved protein signature, independent of TLR2 stimulation. Peptides that did not align to the murine protein library were searched against a *T. gondii* protein library^39^, which identified 86 parasite proteins enriched across the six samples (Figure S1). These included rhoptry protein (ROP)7 and ROP5, which localize to the cytoplasmic face of the vacuole after being secreted into the host cytosol^49^.

As anticipated, several IIGs were highly enriched in PVM samples under ‘IFNγ’ and ‘IFNγ and Pam3CSK4’ stimulation, which reflects the region surrounding all parasites in the sample (Figure 2D). These included IIGP1 (IRGa6), TGTP2 (IRGb6), GBP1, GBP2, GBP5, and GBP7 which have previously been shown to localize to the *T. gondii* vacuole membrane^16, 17, 19^. In addition, GM4951, a predicted IFN-inducible GTPase, and GVIN1, a very large IFN-inducible GTPase, whose function in *T. gondii* restriction has not been explored, were also identified in proximity to the vacuole. The regulatory IRGs, IRGM1, IRGM2, and IGTP (IRGM3) were also identified in the proximity of the parasitophorous vacuole membrane in IFNγ-stimulated conditions. Regulatory IRGMs are thought to be localized to normal self-membranes to block effector IIG attack, although IRGM2 has been observed at the PV in MEF cells^15, 22^. Their enrichment near the vacuole may reflect a sampling of membranes proximal to the vacuole membrane, as the limit of autoSTOMP resolution is larger (∼1um) than the parasite vacuole membrane and intravacuolar network.

This enrichment of IIGs was statistically significant in pairwise comparison between unstimulated PVM samples and IFNγ or IFNγ and Pam3CSK4 treated PVM and GBP2^+^ regions (Figure 2E, PVM). The oligomerization of IRGs and GBPs is integral to their membrane-disruption function^20^, which may explain why the IIG signature was so robust, even in the PVM fraction which reflects a mixture of IIG^+^ and IIG^-^ vacuoles. Consistent with this interpretation, few proteins were significantly differentially enriched when PVM and GBP2^+^ samples were compared within the ‘IFNγ’ or the ‘IFNγ and Pam3CSK4’ stimulated conditions (Figure S2). These data indicate that autoSTOMP is an efficient tool to selectively tag and identify proteins enriched in the vicinity of the PV membrane during resting and immune-stimulated conditions in primary immune cells.

Aside from IIGs, inducible nitric oxide synthase (iNOS, encoded by the gene *Nos2*) was among the most enriched protein in the PVM and GBP2^+^ regions of IFNγ and Pam3CSK4 stimulated condition (Figure 2D, 2Eiv, purple). iNOS was also enriched in the IFNγ-stimulated GBP2^+^ region relative to the unprimed condition, although this was not statistically significant due to variation in the replicates (Figure 2D, 2Eiii). Nitric oxide synthetases (endothelial eNOS, neuronal nNOS, and iNOS) produce nitric oxide (·NO) a membrane-diffusible, radical gas that regulates cell signaling^50^. iNOS is unique in its capacity for transcriptional upregulation in myeloid cells leading to millimolar concentrations of ·NO and conversion to microbicidal reactive species^51^.

### iNOS expression in myeloid cells is necessary for mice to survive acute *T. gondii* infection

An essential role for iNOS in host resistance to *T. gondii* was first determined in 1997 using whole-body knockout mice^52, 53^. To determine if iNOS expression in myeloid cells was necessary to protect the host from *T. gondii* in vivo we crossed mice driving Cre expression from the Lys2 promoter (*LysM^cre/wt^*) onto the *Nos2^fl/fl^* background. Infected *LysM^cre^Nos2^fl/fl^* met euthanasia requirements (Figure 3A, pink), with delayed kinetics compared to whole body knockouts (Figure 3A, violet), exhibiting more extreme weight loss during acute infection than *LysM^wt^Nos2^fl/fl^* controls (Figure 3B, grey).

**Figure 3.**
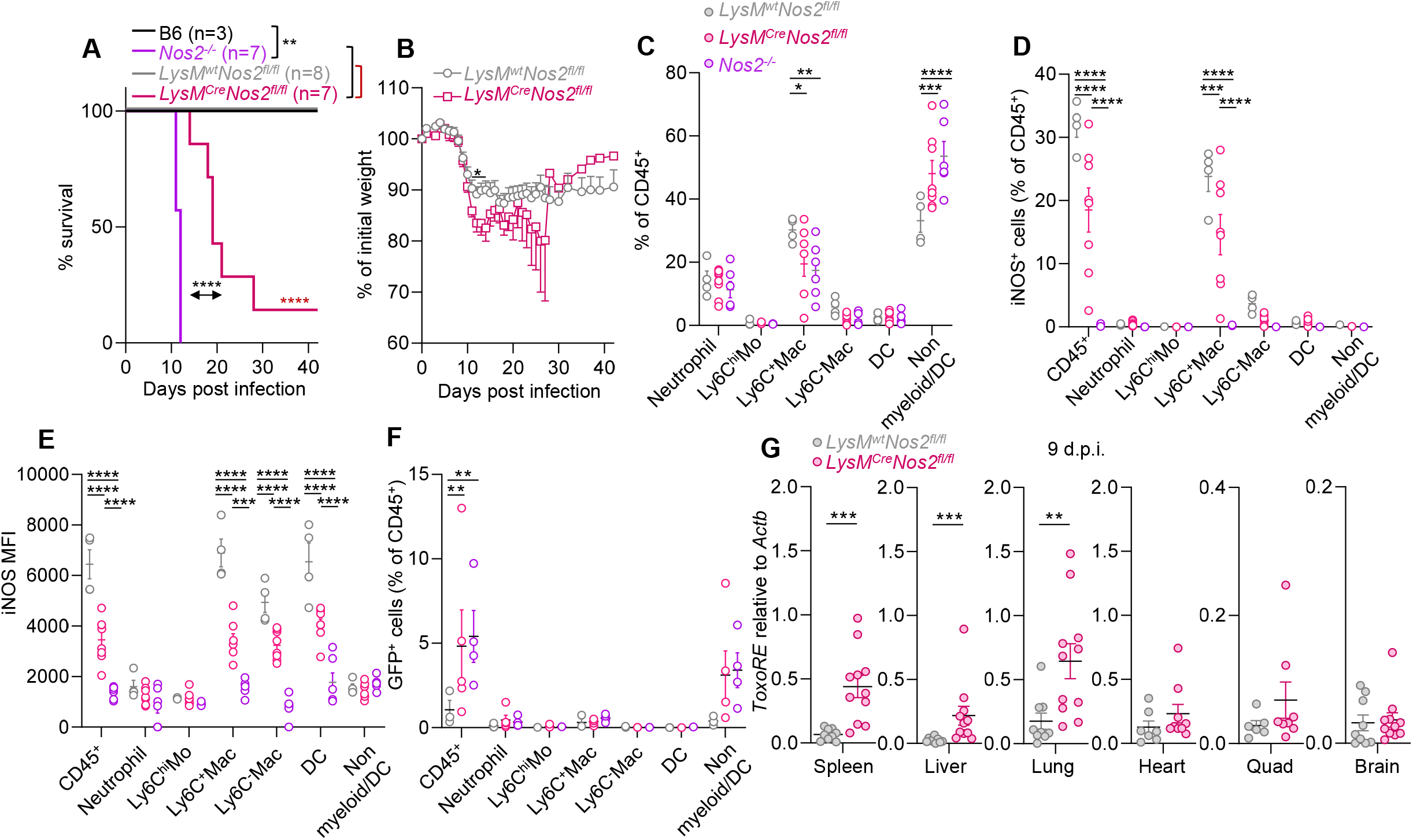
iNOS expression on myeloid cells is necessary for *T. gondii* restriction in vivo and host survival. A-G. Age and sex-matched B6 (black) and *Nos2^-/-^* (violet) mice or *LysM^wt^Nos2^fl/fl^* (grey) and *LysM^Cre^Nos2^fl/fl^* (magenta) littermates were intraperitoneally injected with 5 Me49-GFP-Luc cyst. Euthanasia criteria (**A**, Log-rank test) and weight loss (**B,** student t-test) were monitored for 42 days post-infection (dpi). **C-F**, 8-9 d.p.i. peritoneal exudate was collected and CD45^+^ cells were evaluated by flow cytometry. Neutrophil (CD11b^+^F4/80^-^Ly6G^+^), Ly6C^hi^ monocytes (Ly6C^hi^ Mo, CD11b^+^F4/80^-^Ly6G^-^IA/IE^-^Ly6C^hi^), Ly6C^+^ macrophage (Ly6C^+^ Mac, CD11b^+^F4/80^+^Ly6G^-^IA/IE^+^ Ly6C^+^), Ly6C^-^ macrophage (Ly6C^-^ Mac, CD11b^+^F4/80^+^Ly6G^-^IA/IE^+^ Ly6C^-^), dendritic cells (DC, CD11c^+^IA/IE^+^), non-myeloid or DC (non myeloid/DC, CD11b^-^CD11c^-^Ly6G^-^F4/80^-^) were analyzed from *Nos2^-/-^* (N=6), *LysM^wt^Nos2^fl/fl^* (N=4) and *LysM^Cre^Nos2^fl/fl^* (magenta N=6) animals. **D-E,** For each population, iNOS positive cells were gated (**D**) and iNOS mean fluorescence intensity (MFI) was determined (**E**). **F,** For each population, *Tg-GFP* positive cells were gated. **G**, Parasite burden in the spleen, liver, lung, heart, skeletal muscle (Quadricep, Quad), and brain genomic DNA were analyzed by qPCR using primer probes specific to *ToxoRE* normalized to mouse *Actb,* N=9 *LysM^wt^Nos2^fl/fl^* and N=11 *LysM^Cre^Nos2^fl/fl^*, two tailed Mann-Whitney test. *, p≤0.05; **, p≤0.01; ***, p≤0.001, ****, p ≤0.0001.

To evaluate iNOS-dependent changes in immune infiltrate peritoneal exudate cells (PECs) were isolated for flow cytometry nine days post-infection, before *Nos2^-^*^/-^ mice became moribund. Compared to *LysM^wt^Nos2^fl/fl^* mice, *Nos2^-l-^* and *LysM^cre^Nos2^fl/fl^* mice had a significant reduction in Ly6C^+^ macrophages as well as a compensatory increase in small, panel-negative cells, which were likely lymphocytes (Figure 3C, Figure S3). There was a trend towards fewer Ly6C^-^ macrophage and dendritic cells in iNOS-deficient mice, whereas neutrophils and Ly6C^hi^ inflammatory monocytes were similar across genotypes (Figure 3C). Approximately 30% of CD45^+^ PECs were iNOS-positive in *LysM^wt^Nos2^fl/fl^* mice the majority of which were Ly6C^+^ macrophages (Figure 3D). In *LysM^cre^Nos2^fl/fl^* PECs there was a significant reduction in iNOS^+^Ly6C^+^ macrophages, although this number was highly variable from mouse to mouse compared to the *Nos2*^-/-^ controls. This result is consistent with a study demonstrating that LysM-Cre mice could drive nearly 90% Rosa-fl-STOP-fl-YFP reporter expression in peritoneal macrophages, however, stop codon excision was only 40% efficiency in peripheral blood monocytes^54^. The variation may reflect a scenario where iNOS is upregulated in the myeloid progenitors before the LysM promoter is activated to drive Cre-mediated excision of *Nos2* exons, resulting in residual levels of iNOS proteins in the differentiated cells. Consistent with this model, the mean fluorescence intensity (MFI) of iNOS staining was significantly lower in the iNOS^+^*LysM^cre^Nos2^fl/fl^* Ly6C^+^macrophages, Ly6C^-^ macrophages and dendritic cell populations than *LysM^wt^Nos2^fl/f^*mice (Figure 3E). LysM-cre has been reported to drive gene deletion in neutrophils, however, iNOS expression was not observed in neutrophils, Ly6C^hi^ inflammatory monocytes, or the panel negative cells relative to *Nos2*^-/-^ controls (Figure 3D-E).

Parasite levels are low in the peritoneal cavity by nine days post-infection, nevertheless, there was a significant increase in GFP^+^ (*T. gondii*) CD45^+^ cells in the *Nos2*^-/-^ compared to the *LysM^wt^Nos2^fl/f^* (Figure 3F). *T. gondii* load was significantly higher in the spleen, liver, and lung of *LysM^cre^Nos2^fl/fl^* compared to *LysM^wt^Nos2^fl/lf^* littermates (Figure 3G), sites of dissemination from the peritoneal cavity. There was a trend toward higher parasite levels in the skeletal muscle, heart, and brain, however, this was not significant, which may reflect the early stage of parasite colonization of these tissues. Based on these data we conclude that optimal iNOS expression in the macrophage and monocyte lineages is necessary to control *T. gondii* burden and host survival following intra-peritoneal infection.

### iNOS expression and activity are necessary for chromosome 3 GBP-mediated *T. gondii* restriction

Our experiments demonstrating that iNOS expression in the myeloid cells is necessary for parasite restriction in vivo as well as the observation that iNOS is enriched near GBP2^+^ PVs by autoSTOMP led us to hypothesize that iNOS is necessary for cell-autonomous restriction of *T. gondii* in GBP-targeted vacuoles. Consistent with the literature, IFNγ stimulation positively regulated *Nos2* transcript expression (Figure 4A) and iNOS protein levels (Figure 4B) in a way that was significantly enhanced by TLR stimulation^55^. Notably, *T. gondii* infection was sufficient to synergize with IFNγ to upregulate iNOS, which is likely due to NF-κB phosphorylation by the parasite effectors GRA15^56^, GRA24^57, 58^ and/or parasite ligation of TLRs^7^. By comparison, *Nos1* and *Nos3* transcripts were not detected in BM-MoDCs (Figure S4A).

**Figure 4.**
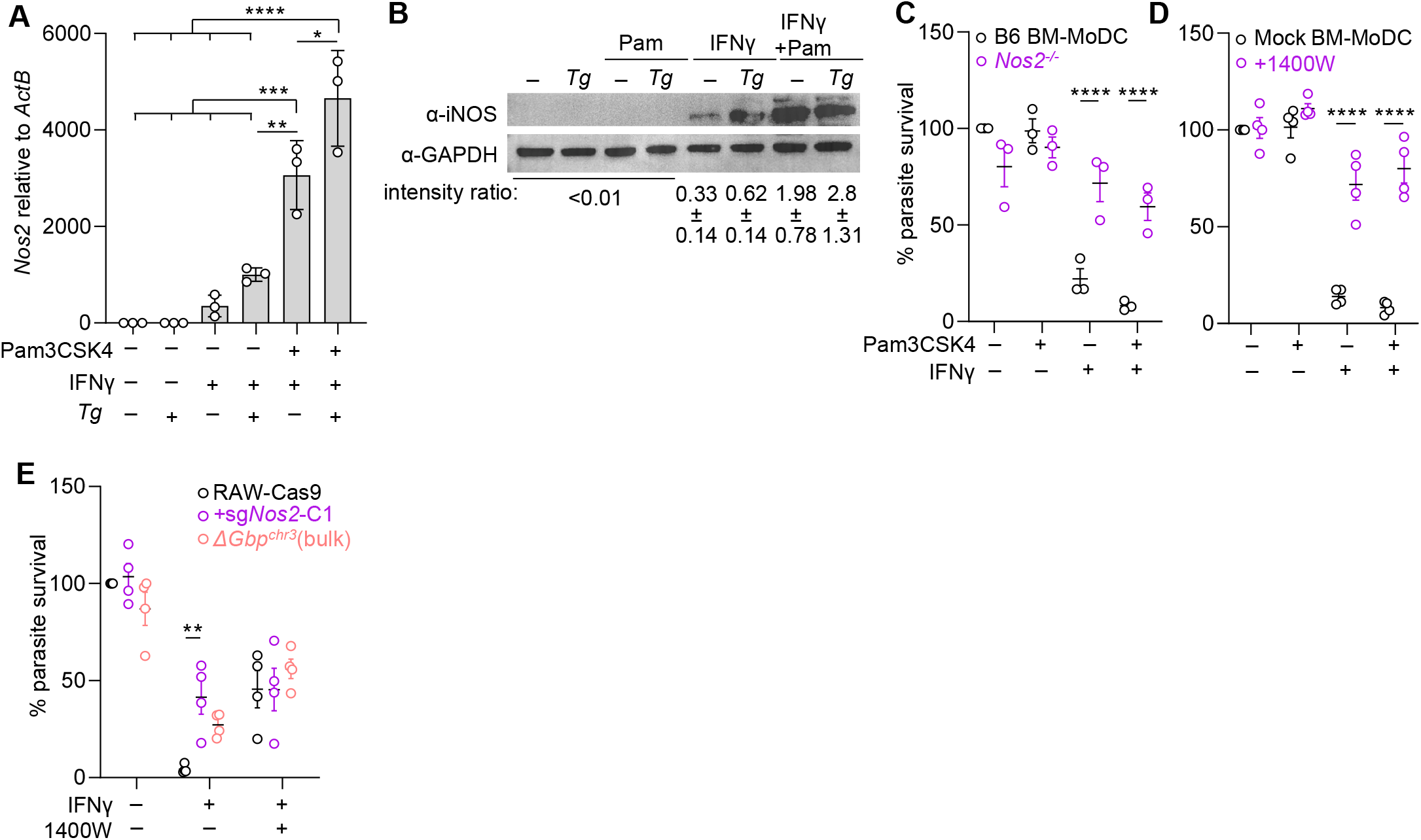
**iNOS is upregulated by IFNγ and *T. gondii* infection and necessary for chromosome 3 GBP-mediated *T. gondii* restriction in BM-MoDCs and RAW 264.7 macrophages. A-D**, BM-MoDCs were stimulated and infected as described in Figure 1. **A**, *Nos2* transcript expression was determined by quantitative RT-PCR at 2 hpi. **B**, iNOS protein levels were detected by Western blot at 14hpi. **C-D**, Parasite survival was measured by luciferase assay 14hpi in B6 (black) or *Nos2^-/-^* (violet) BM-MoDCs (**C**), treated with the iNOS inhibitor 1400W or vehicle control (**D**). **E**, Parental RAW 264.7 macrophages stably expressing Cas9, with a CRISPR-mediated deletion of *Nos2*, or deletion of the chromosome 3 GBP locus *ΔGbp^chr3^* were treated with media or 10ng/ml IFNγ for 20 hours with1400W or vehicle. Cells were infected with Me49GFP Luc at MOI 5 and parasite survival was measured at 14 hpi. Error bars represent Mean±SEM by one-way ANOVA with Tukey post hoc analysis (**A**), 2-way ANOVA with Šidák (**C**, **D**) or Tuckey (**E**) post hoc analysis. *, p≤0.05; **, p≤0.01; ***, p≤0.001; ****, p≤0.0001. N=3 or 4 independent experiments for each panel.

To determine if iNOS was necessary for IFNγ-mediated *T. gondii* restriction, we infected BM-MoDCs from B6 or *Nos2^-/-^* animals. INOS deficiency rescued *T. gondii* growth in BM-MoDCs treated with IFNγ or IFNγ and Pam3CSK4 to levels that were statistically similar to unstimulated BM-MoDCs (Figure 4C). To determine if iNOS activity was necessary for IFNγ-mediated *T. gondii* restriction B6 BM-MoDCs were treated with the selective iNOS inhibitor 1400W (Figure 4D), which rescued parasite growth in IFNγ or IFNγ and Pam3CSK4-stimulated conditions as efficiently as deleting *Nos2* (Figure 4C). In addition, we confirmed that *Nos2*-deletion did not rescue parasite growth by negatively impacting the transcript expression of chromosome 3 GBPs, effector IRGb6 and IRGb10 (Figure S4C), or GBP2 protein levels in response to IFNγ (Figure S4B-C).

To determine if IIGs were necessary for iNOS-mediated parasite restriction we interrupted *Nos2* (RAW*^ΔNos2^*) or deleted the chromosome 3 GBPs (GBP1, -2, -3, -5, and -7, RAW*^ΔGbp-chr3^*) in RAW 264.7 cells using CRISPR/Cas9 (Figure S5A-C). Like BM-MoDCs, iNOS was necessary for optimal parasite restriction in response to IFNγ (Figure 4E, violet). Consistent with previous reports in macrophages, deleting chromosome 3 GBPs rescued parasite growth in IFNγ-treated RAWs to ∼30% of unstimulated levels (Figure 4E, red vs. black)^18^. The RAW*^ΔGbp-Chr3^* deletion was less efficient at rescuing parasite growth than the iNOS deletion (Figure 4E, violet), which is likely due to the functional redundancy of chromosome 5 GBPs and/or IRGs^18^. Importantly, treating IFNγ-stimulated RAW*^ΔGbp-chr3^* cells with 1400W did not enhance parasite growth beyond 1400W treatment alone which is consistent with the interpretation that iNOS and the chromosome 3 GBPs function in the same pathway rather than additively to promote parasite restriction. IFNγ and TLR stimulation induces endosome maturation which could promote parasite clearance independent of IIGs^59^. However, chloroquine, which buffers lysosome acidification, did not rescue parasite growth in IFNγ-treated conditions (Figure S5D). Together these data support a model where chromosome 3 GBPs are necessary for optimal iNOS-mediated parasite clearance.

### iNOS restricts *T. gondii* in IFNγ-stimulated BM-MoDCs through the production of nitric oxide and downstream synthesis of reactive nitrogen species

We next asked which effector functions of iNOS were responsible for *T. gondii* clearance. iNOS converts the substrate L-arginine into nitric oxide (^•^NO) and L-citrulline (Figure 5A)^51^. *T. gondii* is an arginine auxotroph and arginine-limited media impairs tachyzoite growth in fibroblasts^60^. iNOS activation in macrophages has been proposed to limit *T. gondii* growth by depleting free arginine^60, 61^. However, supplementing L-arginine did not rescue parasite growth in wildtype BM-MoDCs stimulated with IFNγ+Pam3CSK4 compared to *Nos2^-/-^* BM-MoDCs (Figure 5B), indicating that iNOS does not mediate parasite clearance via arginine starvation.

**Figure 5.**
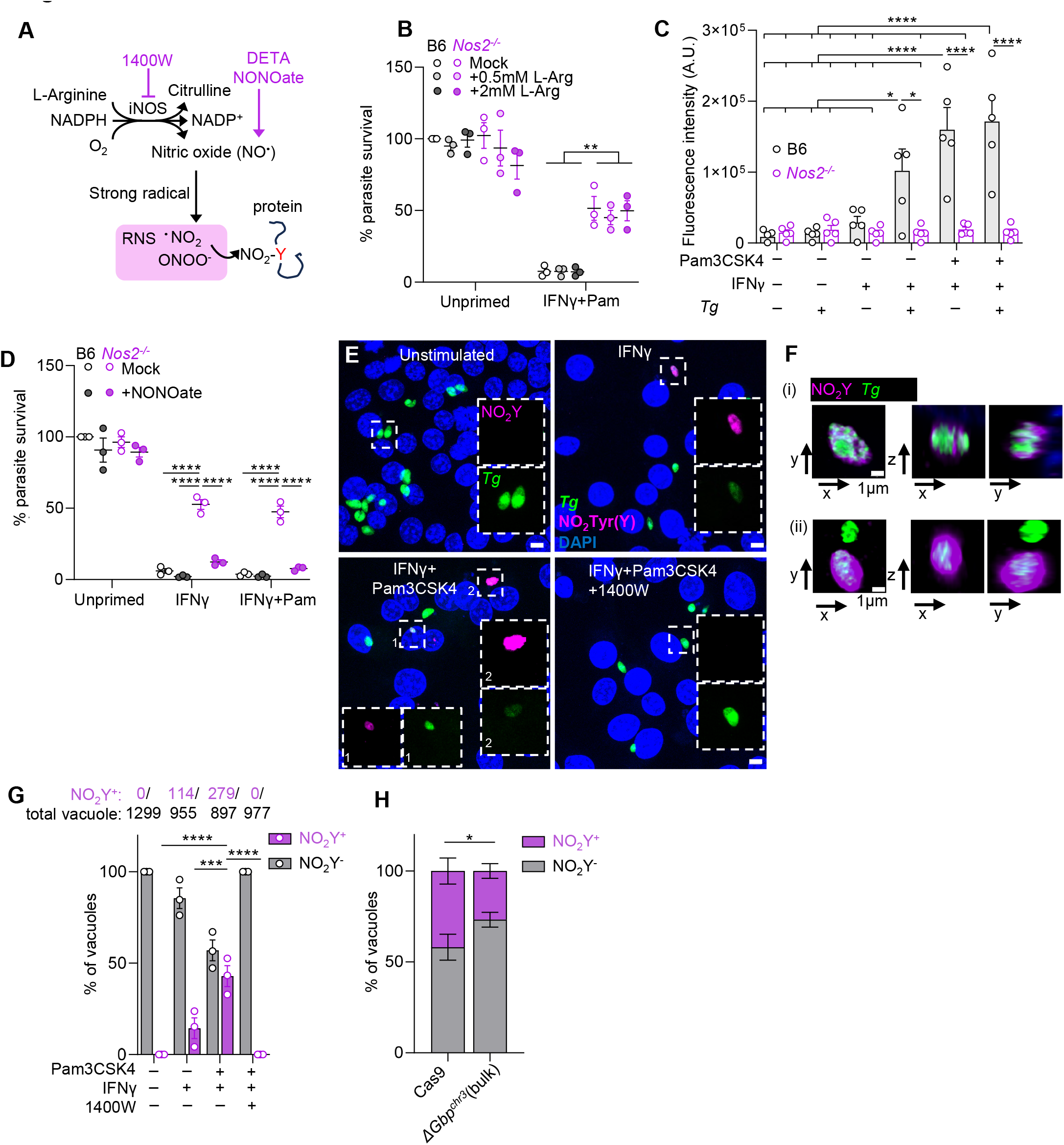
**Reactive nitrogen species are necessary for *T. gondii* restriction, leading to nitration of *T. gondii* vacuoles. A**, Schematic of L-arginine flux through iNOS, to generate NO synthesis, reactive nitrogen species (RNS). **B-D,** BM-MoDCs were stimulated and infected as described in Figure 1. **B,** Parasite growth in normal media or media supplemented with L-arginine one hour before infection was measured by luciferase assay at 14 hpi. **C**, NO production was measured by NO sensitive fluorescent probes at 14hpi. **D**, The NO donor DETA NONOate was added to BM-MoDCs one hour prior to infection, and parasite load was determined by luciferase assay at 14hpi. **E-H**, At 6 hours post-infection, samples were fixed and stained with a nitrotyrosine-specific (NO_2_-Y, magenta) antibody to evaluate co-localization with parasite GFP (**E**), high-resolution imaging showing one plane from z-stacks (**F**). Scale bar, 5 μm (**E**) or 1 μm (**F**). Co-localization was quantified, in the absence or presence of iNOS inhibitor (1400W) (**G**), (**H**) the requirement for chromosome 3 GBPs was evaluated in a similar experiment using RAW cells, N=4 independent experiments. **B-D**, **G**, N=3 unless noted. Mean±SEM, two-way ANOVA with Tukey post hoc test. **H**, RM two-way ANOVA with Šidák post hoc analysis *, p≤0.05; **, p≤0.01; ***, p≤0.001; ****, p≤0.0001.

• NO is a membrane-permeable gas that does not readily react with most biological molecules until converted to reactive nitrogen species (RNS) via reaction with strong radicals^62^ (Figure 5A). RNS, including nitrogen dioxide (^•^NO_2_) and peroxynitrite (ONOO^-^), regulate a range of biological processes by modifying proteins, unsaturated lipids, and nucleic acids^63, 64^. ^•^NO was significantly induced by *T. gondii* infection in IFNγ-stimulated BM-MoDCs or by Pam3CSK4 and IFNγ priming (Figure 5C, grey bars), mirroring the pattern of iNOS protein expression (Figure 4B). As expected, ^•^NO production was entirely dependent on functional *Nos2* (Figure 5C, violet bars). To determine if supplementing exogenous ^•^NO was sufficient to restrict *T. gondii*, we treated B6 or *Nos2^-/-^* BM-MoDCs with the ^•^NO donor DETA NONOate (Figure 5D). DETA NONOate was sufficient to restore IFNγ-mediated *T. gondii* restriction in *Nos2^-/-^* BM-MoDCs. Notably, DETA NONOate did not restrict *T. gondii* in unstimulated cells (Figure 5D) consistent with a model where chromosome 3 GBPs and iNOS cooperate to restrict parasite infection (Figure 4E). Similarly, expressing a doxycycline-inducible allele of human iNOS in RAW cells was not sufficient to restrict *T. gondii* in unstimulated conditions (Figure S6B) even though nitrite (NO_2_^-^) levels were equivalent to IFNγ treatment at 400 ng/mL doxycycline (Figure S6A).

To examine the role of reactive oxygen species (ROS) in *T. gondii* clearance, B6 BM- MoDCs were treated with the antioxidant N-acetylcysteine (Figure S7A, NAC) or the mitochondrial ROS scavenger mitoTEMPO (Figure S7B). Unlike the iNOS inhibitor 1400W, treatment with NAC or mitoTEMPO failed to rescue parasite growth in IFNγ or IFNγ and Pam3CSK4-treated B6 BM- MoDCs. In line with this result, NADPH oxidases were not significantly transcriptionally upregulated in infected RAW264.7 cells (Figure S7C) and phagosome oxidase (PHOX) were not enriched at the vacuole in any IFNγ-stimulated autoSTOMP conditions (Supplementary Table 1). Together, these data are consistent with a primary role for RNS downstream of iNOS in GBP-mediated parasite restriction.

Amino acid nitrosylation (R-NO) and nitration (R-NO_2_) are post-translational modifications that regulate protein-protein associations, protein-membrane interactions, protein turnover, and signaling transduction, for example, by competing for phosphorylation sites^64, 65^. Unlike cysteine nitrosylation, which is extremely labile, tyrosine nitration is stable and until recently was thought to be an irreversible modification (Figure 5A)^63, 66^. To determine if the parasite vacuole was a direct target of RNS, we stained infected RAW 264.7 cells with a nitro-tyrosine (NO_2_Y)-specific antibody (Figure 5E-G). Following IFNγ or IFNγ and Pam3CSK4 stimulation robust NO_2_Y staining co-localized with *T. gondii* GFP signal (Figure 5E, Fi) and, often, the region surrounding the parasite signal (Figure 5Fii). Tyrosine nitration was remarkably selective for the parasite vacuole, as little NO_2_Y staining of the host cell was detected. In addition, tyrosine nitration was vacuole selective as RAW 264.7 cells that had been infected multiple times were observed with NO_2_Y-positive and NO_2_Y-negative vacuoles (Figure 5Fii). Tyrosine nitration was reversed by treatment with the iNOS inhibitor 1400W (Figure 5G). Like our parasite restriction result (Figure 5D), supplementing the nitrogen donor NONOate in unstimulated conditions was not sufficient to induce appreciable vacuole nitration (Figure S7D). This result suggested that IIGs downstream of IFNγ were necessary for vacuole nitration. To test this directly, we evaluated tyrosine nitration in RAW cells deficient in chromosome 3 GBPs (Figure 5H) and found that following IFNγ and Pam3CSK4 treatment, there was a significant reduction in NO_2_Y-positive vacuoles compared to RAW-Cas9 cells. Based on these data we conclude that chromosome 3 GBP recruitment to the vacuole is necessary for optimal RNS-mediated clearance of parasites downstream of iNOS activation and synthesis of NO.

### iNOS is dispensable for GBP2 recruitment to the vacuole and PV membrane ruffling but necessary for intravacuolar network (IVN) membrane collapse and *T. gondii* clearance

IIG localization to the parasite vacuole membrane has been associated with membrane ruffling, permeabilization to cytosolic fluorescent molecules, and *T. gondii* death^16, 17, 20, 27, 33^. To understand how iNOS cooperates with GBPs to mediate parasite clearance, we asked if iNOS functions upstream or downstream of GBP recruitment to the vacuole. RAW-Cas9 cells and RAW*^ΔNos2^* cells had a similar frequency of GBP2^+^ vacuoles by immunofluorescence assay (Figure 6A-Bi), indicating that iNOS was not necessary for GBP2 recruitment. It was immediately apparent that many of the GBP2-positive vacuoles in iNOS-deficient RAW*^ΔNos2^* cells contained morphologically normal, dividing parasites (Figure 6A, arrowhead). The number of parasites per GBP2-positive vacuole was significantly increased in RAW*^ΔNos2^* compared to RAW-Cas9 cells (Figure 6Bii). However, the number of parasites per GBP2-negative vacuole was similar, indicating that parasites were not simply growing more efficiently in RAW*^ΔNos2^* than RAW-Cas9 cells (Figure 6Bii). These data indicate that IIG recruitment is not sufficient to optimally restrict parasite growth in the absence of iNOS, supporting a model of cooperative clearance downstream of IIG recruitment.

**Figure 6.**
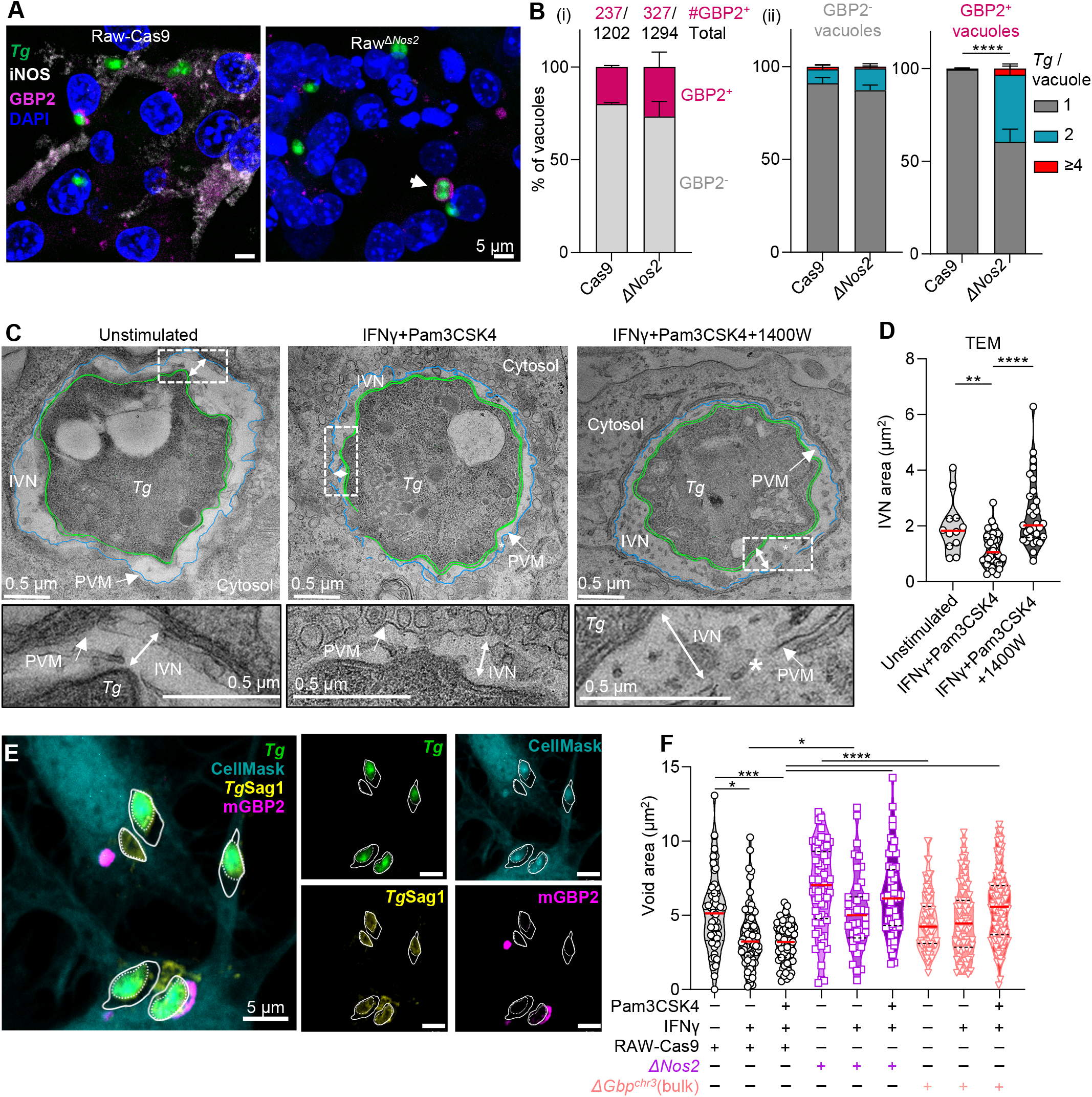
**iNOS leads to intravacuolar network collapse and is necessary to restrict parasite growth in GBP-targeted vacuoles. A-B**, RAW-Cas9 or RAW*^ΔNos2^* cells were infected with Me49-GFP-Luc as described in Figure 4. 6hpi, samples were fixed and stained with antibodies specific to GBP2 (magenta) and iNOS (white) (**A**), arrow indicates dividing parasites in GBP2-positive vacuole. **B**, The frequency of GBP2-targeting (i), and the number of parasites in GBP2-negative (grey) or GBP2-positive (magenta) vacuoles was quantified (ii). N=3 independent experiments, Mean±SEM, two-way ANOVA with Šidák post hoc analysis. **C-D,** Vacuole ultrastructure was evaluated by transmission electron microscopy in RAW-Cas9 cells that were unstimulated, treated with IFNγ and Pam3CSK4 alone or with 1400W then infected for 6hours. Insets show the intravacuolar network between the parasite plasma membrane (green line, *Tg*) and the parasite vacuole membrane (blue line, PVM), * indicates breaks in the PVM (**C**). **D**, Samples were blinded, and the intravacuolar network area (IVN) quantified. One representative experiment, Kruskal-Wallis test with Dunn’s post hoc analysis. **E-F**, To measure vacuole integrity, RAW-Cas9 (open circle), RAW*^ΔNos2^* (open square) or RAW*^ΔGbp-chr3^* (open triangle) cells were infected with Me49-GFP-Luc as described in Figure 4. 6hpi, samples were fixed and stained with CellMask (aqua) and an antibody specific to *Tg*SAG1 (yellow). The IVN area between the cell mask signal (solid white line) and the parasite GFP and/or SAG1 signal (dotted line) was quantified (**F**). Pooled data of 2 independent experiments showing measurements of individual vacuoles, Kruskal-Wallis test with Dunn’s post hoc analysis. *, p≤0.05; **, p≤0.01; ***, p≤0.001; ****, p≤0.0001.

To understand how iNOS impacts parasite vacuole structure we infected RAW-Cas9 cells that were not stimulated, stimulated with IFNγ and Pam3CSK4 alone, or with iNOS inhibitor using transmitted electron microscopy (TEM) (Figure 6C). The intra-vacuolar network (IVN) is a network of host-derived lipid nanotubes and parasite effectors that connect parasites to the PV membrane and to each other. Consistent with previous reports^17, 25, 27, 33^, IFNγ and Pam3CSK4 treatment led to membrane ruffling, discontinuity, and a significant reduction in the IVN area (Figure 6C-D). Unexpectedly, blocking iNOS activity rescued the IVN volume (Figure 6C-D, double arrow, and inset) although morphological changes to the parasite vacuole membrane were still observed (Figure 6C, asterisk). Notably, the loss of PV membrane continuity was also observed in 1400W- treated vacuoles containing two parasites (Figure S8), suggesting that parasite viability was intact. Together, these data are consistent with a model where IIGs disrupt the PV membrane independent of iNOS activity, however, disruption of the membrane is not sufficient for efficient parasite killing in the absence of iNOS-mediated IVN collapse.

To understand the requirement of iNOS and the chromosome 3 GBPs on vacuole integrity, we labeled the host cell cytosol with CellMask, blinded the samples, then evaluated the area of the CellMask-negative IVN ‘void’ surrounding each parasite (Figure 6E). In RAW-Cas9 cells, IFNγ treatment led to a significant reduction in the void area between the *T. gondii* and CellMask (Figure 6F, black). Similar to the TEM results (Figure 6D), the CellMask-negative void was rescued in RAW*^ΔNos2^* cells treated with IFNγ or IFNγ and Pam3CSK4 (Figure 6F, violet). RAW*^ΔGbp-chr3^* cells also had a significant rescue of the CellMask void area surrounding the parasites, (Figure 6F, red). Together these data are consistent with the conclusion that GBP localization to the vacuole and ruffling of the PV membrane is necessary but not sufficient for parasite clearance in mouse macrophage infection. Optimal parasite killing requires transcriptional upregulation of iNOS and synthesis of ^•^NO and RNS which modify the PV and collapse the intravacuolar network leading to optimal parasite clearance in cooperation with the chromosome 3 GBPs (Figure 7).

**Figure 7.**
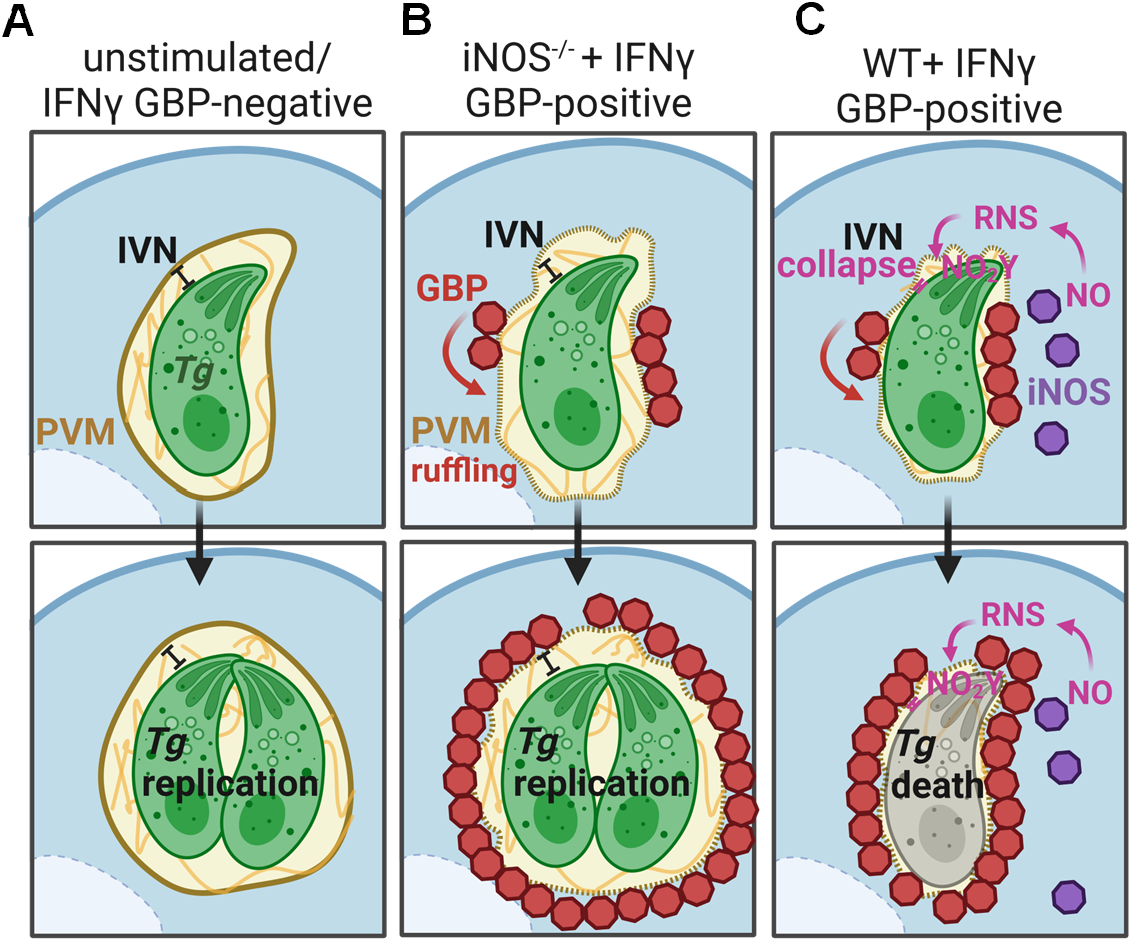
Model of the cooperative iNOS and chromosome 3 GBP-mediated *T. gondii* restriction in murine myeloid cells. A. In unstimulated myeloid cells or IFNγ-stimulated conditions where the PV is not targeted by GBPs parasites are replication competent. **B,** In iNOS- deficient macrophages, GBP recruitment and parasite vacuole membrane (PVM) ruffling occurs, but parasites are replication-competent. **C,** In WT macrophages, iNOS-dependent synthesis of nitric oxide (NO) and reactive nitrogen species (RNS) leads to tyrosine nitration of the PV, collapse of the intravacuolar network, and death of parasites in GBP targeted vacuoles

## Discussion

Here we have shown that iNOS expression in murine macrophages and dendritic cells is necessary for efficient, vacuole-autonomous parasite restriction by GBPs. While iNOS has been implicated in *T. gondii* restriction since the 1990s, the precise mechanism of parasite clearance or a direct relationship with IRG and GBP-mediated killing has not been rigorously assessed. Early reports evaluating the role of iNOS in *T. gondii* infection^61^ could not discriminate between L-arginine deprivation and RNS production as the mechanism of parasite control due to the use of reversible L-arginine analogs to inhibit iNOS^67^. Our data exclude L-arginine deprivation and implicate ^•^NO synthesis as the critical pathway for killing, leading to vacuole nitration in a manner that depends on the chromosome 3 GBPs. Our data align with a published result from the Yamamoto lab showing that iNOS inhibitor treatment increased the number of parasites per vacuole in peritoneal macrophages from WT mice, and this rescue was less penetrant in macrophages from chromosome 3 GBP-deficient mice^18^. However, since infection rate was not dependent on iNOS inhibition, this relationship was not explored further. In addition, skeletal muscle cells treated with IFNγ and TNF have been shown to recruit IRGb6 to the vacuole and produce nitrite in response to Type I and Type II parasite infection, however, a direct relationship between these effector mechanisms was not tested^68^. Perhaps the most critical evidence supporting the importance of iNOS in *T. gondii* clearance is the observation that parasites have evolved strategies to inhibit the ^•^NO production^69, 70^ and promote the proteasomal degradation of iNOS^69^. In 2007 the Knoll lab found that *T. gondii* patatin-like protein (*Tg*PL1) allowed *Tg* to limit nitrite synthesis in LPS and IFNγ-stimulated macrophages and protected parasites from degradation^71, 72^, however, the precise mechanism by which *Tg*PL1 interrupts the iNOS/^•^NO axis is not clear.

Our work significantly expands on these previous studies by establishing that the PV is robustly and selectively modified by reactive nitrogen species (Figure 5D). The high levels of ^•^NO flux observed in these studies are expected to lead to dynamic nitration and nitrosylation of lipid, protein and/or nucleic acid substrates^63^. Although we could not detect nitrated nucleic acids in the parasite using 8-nitroguanine specific antibodies (data not shown), it is likely that other targets of RNS, including reactive cysteines and unsaturated lipids^63, 64^, also participate in IVN collapse. For example, unsaturated phospholipid tails in lipid bilayers can serve as nitrogen sinks that target membrane integral proteins for nitration and nitro-lipidation^73^. In addition, lipid nitration has been proposed to regulate lipid membrane dynamics using synthetic micelles and modeling experiments^63, 74^, although this has not been shown in cells. Reliable identification of nitrated lipid species is challenging and there are currently no tools to evaluate the sub-cellular location of modified lipids. Until recently, tyrosine nitration was posited to be irreversible^63, 66^, which may be why this modification was readily detected by immunofluorescence assay in our study. While it is formally possible that a specific NO_2_Y-modified substrate(s) regulate the processes of IVN collapse and parasite clearance in partnership with GBP targeting, we favor a model where functional redundancy of nitrated/nitrosylated proteins and or lipid targets mediate this biology^63^. Future mass spectrometry experiments will be required to identify individual targets and test the role of these modifications on a case-by-case basis.

The IVN (also called the membranous nanotubular network or, in *Plasmodium* the tubovesicular network) is a series of host-derived lipid channels connecting the PV membrane to the parasites and parasites to each^3, 41^. Initially, the IVN was thought to mediate nutrient acquisition from the host, however, emerging data suggest that the IVN may provide structural support for the parasite and is necessary for synchronous division^75, 76^. Many parasite rhoptry and dense granule proteins traffic to the IVN although their roles in this compartment are not entirely clear. It is possible that RNS-modification of parasite proteins in the IVN is leading to growth arrest. Although a series of rhoptry proteins were identified in our autoSTOMP data set, we did not identify IVN-associated dense granule proteins (Figure S1). This may be related to the low coverage of this technique (∼1.0% of 8,315 *T. gondii* proteins annotated in UniProt) and/or the low abundance of individual GRA proteins rather than their true absence from the GBP2-targeted region. Future studies using autoSTOMP to target the NO_2_Y-positive vacuolar regions may identify host and/or parasite proteins targeted by RNS as well as mediators of downstream vacuole clearance.

Discovering this role for iNOS has also revealed that GBP recruitment is necessary but not sufficient for IFNγ-dependent parasite clearance in murine macrophages, as previously thought. Our study is consistent with previous TEM studies showing that PV membrane ‘ruffling’ in wild-type macrophages is limited by deletion of the chromosome 3 GBPs^18^ or GBP1^27^. However, our data indicate that these morphological changes to the PV membrane may not be sufficient for the diffuse IVN protein staining^17^, PV membrane permeability and dead parasite phenotype as iNOS inhibition leads to enrichment of GBP2 positive vacuoles containing 2 or more parasites ^16, 33^. Our data support a dual requirement for the chromosome 3 GBPs in identifying and deforming the PV membrane and RNS modification of the IVN for parasite clearance. This 2-hit model is conceptually consistent with the mechanism discovered by Gaudet *et al.* where human GBP1 localization to bacterial membranes must partner with an executioner molecule (apolipoprotein (APO) L3) to mediate bacterial killing^35^. The role of APOL3 and iNOS in *T. gondii* clearance from human macrophages remains to be explored. In comparison to the human system where GBP1 is necessary and sufficient for vacuole disruption^33, 34^, in mice IFNγ upregulates the expression of many, partially redundant of IIGs. IRGa6^26^, IRGb6^25^, IRGd^21^, GBP1^27^, GBP2^28^, and GBP7^29^ have been shown to be necessary for optimal parasite restriction while others, including GBP3, GBP5, and GBP6, can be recruited to the vacuoles already decorated with GBPs^20^. The precise function of each IIGs on the PV membrane is not fully understood, and future studies will be necessary to discern which IIGs are necessary for iNOS-mediated IVN collapse.

Finally, our study is the first to formally demonstrate that iNOS expression in myeloid cells is necessary to control *T. gondii* infection in vivo. Previous studies showed that mice lacking *Nos2* succumb to *T. gondii* by 4 weeks post-infection following IP and oral infection^52, 53^. Lethality was initially attributed to defective parasite restriction in the brain^53^, however, subsequent studies demonstrated that iNOS is necessary to control hepatic parasite burden^52^, and iNOS expression in irradiation-sensitive cells was largely responsible for *T. gondii* protection^77^. Interestingly, *Nos2^-/-^* animals had fewer necrotic lesions in the liver and intestine than wild-type animals, indicating iNOS-mediated parasite clearance comes at a cost of inflammatory damage-to-self^52^. In our studies, not only did *Nos2^-/-^* mice fail to control parasite burden in the peritoneum, liver, spleen, and lung, they met euthanasia requirements by 10 days post-infection (Figure 3), which was more rapid than previous reports. While it is possible that this discrepancy is due to differences in the gene interruption (our study used deletion of *Nos2* exons 12 and 13^78^ previous studies used deletion of *Nos2* exons 1-4^52, 53^), the 129Sv mice used to generate the knockout alleles are notably more resistant to acute infection than C57BL/6 mice^79^. Thus, the limited number of backcrosses used in previous studies (F2 crosses^53^ or 5 backcrosses to C57BL/6^52^ compared to the 11 backcrosses to C57BL/6 mice used here^78^) likely accounts for the delayed lethality. Taken together, this study establishes that iNOS is necessary for efficient, cell-autonomous parasite clearance by the chromosome 3 GBPs in murine myeloid cells.

## Limitations of Study

A limitation of this study is that we are not able to determine if the IFNγ-dependent, chromosome 3 GBP-independent function of iNOS is entirely dependent on the recruitment of chromosome 5 GBPs and/or IRGs^14^. It is formally possible that some vacuoles are sensitive to nitration and IVN collapse in the absence of IIG recruitment, although experiments using peroxynitrite treatment and lysosome acidification inhibitors did not support this alternative model. Unfortunately, genetic tools to eliminate all GBPs and IRGs from murine cells are not available to test this hypothesis. An additional limitation is that the ∼1µm biotinylation region used in AutoSTOMP balances the needs to cross-link sufficient proteins for LC-MS, and cross-linking time (both limiting factor) with the optimal spatial resolution of targeting^39^. This excitation region is significantly larger than the PV membrane. As a result, the proteins identified in these experiments require post-validation to confirm localization to the PV or near neighbor regions of the host cell or vacuole.

## Author contributions

Conceptualization: X.Y.Z., S.E.E.; Research Design: X.Y.Z., S.E.E.; Investigation: X.Y.Z., S.L.L., J.U.A., B.Y., N.K.H.; Data analysis: X.Y.Z., S.L.L.; Image quantification: X.Y.Z., S.L.L., J.S.; Writing -- Draft and Editing: X.Y.Z., S.E.E.; Supervision: S.E.E.

## Declaration of interests

The authors declare no competing interests.

## Inclusion and diversity

We support inclusive, diverse, and equitable conduct of research. Citations in this manuscript were selected to reflect scientific contribution as well as inclusive gender and geographical diversity, whenever possible.

## Methods

### Animals and husbandry

C57BL/6 (Jax #:000664), *Nos2^-/-^* (B6.129P2-*Nos2^tm1Lau^*/J, Jax #:002609), and CBA/J (Jax #:000656) mice were purchased from the Jackson Laboratory. *LysM^cre^Nos2^fl/fl^* mice(Vilela et al. 2022) were a gift from Drs. André Marette from Laval University and Dr. Frederick H. Epstein from University of Virginia. Mice were bred and housed in accordance with the University of Virginia Institutional Animal Care and Use Committee, Association for Assessment and Accreditation of Laboratory Animal Care, and Institutional Animal Care and Use Committee Protocol 4107-12-21.

### Cell culture

Bone marrow was isolated according to previous publications^80, 81^. Bone marrow was seeded at 1x10^6 in RPMI (Gibco,11875119)+10%FBS containing 20ng/mL of recombinant GM-CSF (PreproTech, 315-03), fresh differentiation media was added on day 3 and BM-MoDCs were plated for infection on day 7. Loosely attached and adherent cells were collected, counted using trypan blue on the Countess cell counter (Thermo Fisher Scientific), and seeded on tissue culture treated 96-well (Corning 3596) or 24-well plates (Fablab, FL7123). RAW264.7 cells were purchased from ATCC (cat. # TIB71) and maintained according to the ATCC protocol. Plated cells were rested at least 12 hours before stimulation.

### Parasite infection in vitro

Type II Me49 parasites expressing GFP and luciferase (Me49GLuc) were maintained for up to 10 serial passages on confluent, primary human foreskin fibroblasts (HFFs) in 3 mL of DMEM (Thermo Fisher Scientific, 11965118) +10% FBS. Intracellular parasites were released by scraping and passage through 22g blunt end needles (Instech Laboratories, LS22/6S). The parasite suspension was centrifuged at 450 rpm for 3 min to remove large debris and then washed twice by centrifuging at 1250 rpm for 6 min each. The pellet was resuspended in DMEM+10%FBS and counted on the hemocytometer. Parasites were then diluted to the indicated multiplicity of infection (MOI) so that 5 uL of suspension was spiked in every well for infection.

For BM-MoDC infections, cells were seeded at 1x10^6^ cell/mL, and media was changed to OptiMEM (Gibco, 31985070) +1% FBS containing 70 ng/mL of mIFNγ (R&D, 485MI100CF) 20 hours before infection. 3 hours before infection Pam3CSK4 (Invivogen, tlrl-pms) was spiked into the wells at a final of 100 ng/mL. Cells were then infected with Me49GLuc at MOI 1.5.

For infection with RAW 264.7 cells, cells were seeded at 1.5x10^5^ cell/mL in DMEM+10%FBS and media was changed to DMEM + 10%FBS containing 10 ng/mL of mIFNγ 24 hours before infection and at the last 3 hours, Pam3CSK4 was spiked in to a final of 100 ng/mL. Cells were then infected with Me49GLuc at MOI 5.

At 14 hours post infection, parasite burden was measured using Steady-Luc Firefly HTS Assay Kit (Biotium, 30028-L2) according to manufacturer’s instructions. The parasite burden was calculated using relative light units (RLU) read from Cytation 5 plate reader (BioTek).

The following inhibitors were added 1 hour before infection in this study: 1400W-HCl (100μM, Selleck Chemicals, S8337), L-Arginine (0.5 or 2mM, MP Biomedicals, 0219462625), DETA NONOate (300μM, Cayman Chemical Company, 82120), N-aceyl-L-Cysteine amide (1mM, Cayman Chemical Company, 25866), mitoTEMPO (200 or 500μM, Sigma, SML0737-5MG). Chloroquine (100μM, Thermo Fisher Scientific, 455240250) was added 1 hour after infection to avoid impacting parasite invasion.

### Immunofluorescence staining and imaging

BM-MoDCs or RAW 264.7 cells were seeded on poly-d-lysine (MP Biomedicals, 0215017550) coated coverslips (Harvard Apparatus, 64-0712) in 24-well plates stimulated and infected as described above. At the time of collection, media was aspirated and cells were fixed by 4% PFA (Electron Microscopy Sciences, 15710) for 15 min at room temperature. Samples were washed twice with PBS, permeabilized with PBS+1% Triton X-100 (Fisher) for 30 min, and blocked with PBS+5% BSA (Fisher) or serum of the host of secondary antibodies (Jackson ImmunoResearch). Coverslips were incubated with primary antibodies overnight at 4°C, followed by 3 washes with PBS, and incubation with fluorescent secondary antibodies. After 3 washes and DAPI staining (Thermo Fisher Scientific, D1306), coverslips were mounted in the ProLong Gold Antifade Mountant (Thermo Fisher Scientific, P36930) or Vectashield Mounting Medium (Vector Laboratories, H-1000-10). Coverslips were imaged on Zeiss LSM 880 (Carl Zeiss) using 40x (Plan-Apochromat NA1.3, Oil DIC M27) or 63x (Plan-Apochromat NA1.4, Oil DIC M27) lenses.

Primary antibodies used: ⍺-GBP2 (Proteintech, 11854-1-AP, 1:500 dilution), ⍺-*Toxoplasma* polyclonal (Thermo Fisher Scientific, PA1-7253, 1:500 dilution), ⍺-*Tg*SAG1 (Thermo Fisher Scientific, MA518268, 1:100 dilution), ⍺-iNOS (BD Biosciences, 610328, 1:500 dilution), ⍺- Ubiquitin (Enzo Life Sciences, BML-PW8810-0100, 1:500 dilution), ⍺-nitrotyrosine (EMD Millipore, 06-284, 1:200 dilution). Secondary antibodies used: Goat anti-Rabbit IgG (H+L) Highly Cross-Adsorbed Secondary Antibody used (at 1:500 dilution), Alexa Fluor 594 (Thermo Fisher Scientific, A11037); AffiniPure Donkey Anti Mouse IgG (H+L), Alexa Fluor 594 (Jackson ImmunoResearch, 715-585-150); Donkey anti Rabbit IgG (H+L) Highly Cross Adsorbed Secondary Antibody, Alexa Fluor 647 (Thermo Fisher Scientific, A31573).

### autoSTOMP procedure and LC-MS

AutoSTOMP was performed as published previously^39, 40^. Briefly, B6 BM-MoDCs were seeded on 18 mm coverslips (Thomas Scientific, 1217N81) at 1.35 x 10^6^ cells per coverslip in 12 well plates. Cells were treated with RPMI + 10%FBS containing 70 ng/mL of mIFNγ 16 hours before the infection and at the last 3 hours, Pam3CSK4 was added to a final concentration of 100 ng/mL.

Cells were then infected with Me49GLuc at MOI 1.5. Samples were harvested at 2 h.p.i by fixing in cold absolute methanol (Fisher) for 20 min on ice and stained following the IF staining protocol including avidin/biotin blocking (Vector Laboratories, SP-2001) following the manufacturer’s instructions. Slides were stored at -30 and mounted immediately prior to autoSTOMP imaging in 1 mM biotin-dPEG3-benzophenone (Quanta BioDesign, Biotin-BP) in 50:50 (v/v) dimethyl sulfoxide (DMSO)/water. Slides were subjected to autoSTOMP procedure on Zeiss LSM 880 microscope (Carl Zeiss) with a Chameleon multiphoton light source (Coherent) and a 25x oil immersion lens (LD LCI Plan-Apochromat 25×/0.81 mm Korr DIC M27). Regions of interest were defined as the 1 pixel region surrounding the *Toxoplasma* signal was defined at the MAP or the region of GBP2+ staining overlapping or immediately adjacent to the *Toxoplasma* signal was defined as the MAP. A detailed protocol and source codes are available at https://github.com/boris2008/Sikulixautomates-a-workflow-performed-in-multiple-software-platforms-in-Windows.

After autoSTOMP crosslinking samples were washed, dissociated from the coverslips in 8 M urea lysis buffer (100 mM NaCl, 25 mM Tris, 2% SDS, 0.1% tween 20, 2 mM EDTA, 0.2 mM PMSF, and 1x Roche cOmpleteProtease inhibitor). Lysates were treated with benzonase (Sigma, E1014- 25KU) and RNase (Sigma, 10109142001) to reduce viscosity then subjected to affinity purification using Pierce Streptavidin Magnetic Beads (Thermo, 88817). Enriched proteins were eluted by boiling at 96 °C for 5 min in Laemmli SDS buffer (10µM DTT, 0.0005% Bromophenol blue, 10% Glycerol, 2% SDS, 63 mM Tris-HCl pH 6.8) and ran on a pre-cast Novex Tris-Glycine Mini Protein gel (Thermo Fisher Scientific, XP04125BOX) at 70 V for 12 min. 1 cm gel fragments were cut from each lane and submitted to University of Virginia Biomolecular Analysis Facility for mass spectroscopy analysis. The samples were run on a Thermo Orbitrap Exploris 480 mass spectrometer system with an Easy Spray ion source connected to a Thermo 75 μm × 15 cm C18 Easy Spray column (trap column first). Mass spectra data were analyzed using MaxQuant (versions 1.6.15.0) following the published pipeline and maxLFQ data were analyzed in Perseus (version 2.0.3.1) using Student’s t test (permutation-based FDR)^39, 40^. Plots were generated using R (version 4.2.0) and GraphPad Prism 9.

Raw proteomics data sets associated with this study are freely available for download on PRIDE https://www.ebi.ac.uk/pride/ accession number: PXD027716

### Mouse infection

Age and sex-matched animals were infected between 12-18 weeks old. 2 weeks before infection, dirty bedding was mixed to normalize the microbiota. Me49GLuc cysts were passed in vivo ib CBA/J mice to harvest cysts from brain tissue as previously published^82^. The brain lysate was 1:10 diluted in the PBS and stained with rhodamine-labeled dolichos biflorus agglutinin (DBA-red, Vector Laboratories, RL-1032-2). GFP and DBA-red double-positive cysts were counted and diluted to 5 cysts per 200 uL of PBS. Mice were injected with 200 uL of cyst solution and were monitored using a humane endpoint scoring system based on weight loss, posture, appearance, and activity. Moribound mice were euthanized according to the ACUC protocol.

### Flow cytometry

Cells in the peritoneal cavity were isolated by lavage with 10 mL of cold PBS. Cell number was determined by Countess hemocytometry (Thermo Fisher Scientific). 1.2x10^6^ total cells were isolated for FACS staining in a round bottom 96-well plate. Cells were pelleted down at 1,500 rpm at 4°C for 5 min and cell pellets were resuspended in 100 uL FACS buffer (PBS + 4% FBS +0.5 mM EDTA) containing Fc block (BioLegend, 101302, 1:200). After 15 min incubation at 4°C, surface markers were stained with following antibodies at 1:200 dilution at 4°C for 30 min: CD11b- Pacific Blue (BioLegend, 101224), CD45-BV650 (BioLegend, 103151), CD11c-BV711 (BioLegend, 117349), Ly6C-PerCP/Cy5.5 (BioLegend, 128012), F4/80-PE/Cy7 (BioLegend, 123113), I-A/I-E-AF647 ((BioLegend, 107618), Ly6G-APC/Cy7 (BioLegend, 127624). Staining was stopped by 2 washes in FACS buffer, fixation, and permeabilized using BD Cytofix/Cytoperm fixation/permeabilization kit (BD Biosciences, 554714). Intracellular iNOS was stained with iNOS- AF594 (BioLegend, 696803, 1:200) in the FACS buffer for 1 hour at room temperature in the dark. Cells were then washed twice using BD Perm/Wash buffer and resuspended in 200 uL of FACS buffer. Single stains were prepared using UltraCom eBeads (Thermo Fisher Scientific, 01-2222- 42) as well as fluorescence minus one (FMO) controls. Samples were run on an Attune Flow Cytometer (Thermo Fisher Scientific) equipped with 405 nm, 488 nm, 561 nm, 637 nm lasers, and 14 detector channels. Data were analyzed using FlowJo (v10.8) and gated using FMOs.

### Genomic DNA preparation from tissue

Tissues were dissected and flash-frozen on liquid nitrogen. 1 µL of UltraPure water (Thermo Fisher Scientific, 10977015) was added per 1 mg of tissue, then bead beaten (Qiagen, 69989) using a Qiagen Tissuelyzer (Qiagen) for 3 min at 25 Hz. 20 µL of tissue lysates were taken for DNA isolation using DNeasy Blood & Tissue Kit (Qiagen, 69506) according to the manufacturer’s instruction. Eluted DNA was quantified on the NanoDrop One (Thermo Fisher Scientific) and diluted to the same concentration. 50 ng of total DNA from each sample was used for quantitative PCR (qPCR).

### Quantitative PCR

*T. gondii* burden in the tissue was measured by qPCR of *T. gondii* 529-bp repeat element (RE) compared with mouse β-actin as described previously^83^ using Taqman probes (Thermo Fisher Scientific). Water or uninfected animals were used as a negative control. Data were analyzed using the ΔCt method. The following TaqMan primer/probes were used: 529-bp RE forward, 5′- CACAGAAGGGACAGAAGTCGAA-3′ and reverse, 5′-CAGTCCTGATATCTCTCCTCCAAGA-3′; probe: 5′-CTACAGACGCGATGCC-3 (Integrated DNA Technologies); mouse β-actin: Mm02619580_g (Thermo Fisher Scientific).

For tissue culture experiments, total RNAs were isolated using Quick-RNA Miniprep Kit (Zymo Research, R1055). RNA concentration and quality were assessed by NanoDrop (Thermo Fisher Scientific). 40 ng of total RNA was used for cDNA synthesis using Accuris qMax cDNA Synthesis Kit (Accuris Instruments, PR2100-C-250). Gene expression levels were measured by reverse transcription quantitative PCR (RT-PCR) using PowerUp SYBR Green PCR Master Mix (Thermo Fisher Scientific, A25742). Primers are listed in the Supplementary Table S3 and ΔCt was calculated relative to *Actb* gene while relative gene expression levels were quantified using ΔΔCt method normalizing to unstimulated groups. No RT control was included as a negative control. All of the above procedures were carried out according to the manufacturer’s instructions.

### Western blot

At the time of collection, media was removed, and cell lysates were prepared by directly adding Laemmli SDS buffer into the well. Protein lysates were transferred out of the plate, sonicated twice on ice, and denatured at 95°C for 5 min. Lysates were loaded on 10% home-made SDS-PAGE gel and transferred onto a 0.45 µm PVDF membrane (Millipore, IPFL00010) using TransBlot Turbo System (Bio-Rad). The membranes were blocked in TBST containing 5% skim milk at room temperature for 1 hour, followed by primary antibody incubation at 4°C overnight. After three TBST washes, membranes were incubated in TBST containing HRP-conjugated secondary antibodies at room temperature for 1 hour. Luminata Forte Western HRP substrate (Millipore, WBLUF0100) and ChemiDoc Imager (Bio-Rad) were used to develop the membrane and acquire images respectively. Volumetric intensities were quantified using Image Lab software (Bio-Rad, version 6.0.0). Primary antibodies used: ⍺-GBP2 (Proteintech, 11854-1-AP, 1:1000 dilution), ⍺-iNOS (BD Biosciences, 610328, 1:1000 dilution), ⍺-GAPDH (CST, 5174, 1:1000 dilution). Secondary antibodies used: Peroxidase AffiniPure Goat Anti-Mouse IgG (Jackson ImmunoResearch, 115-035-003, 1:10,000 dilution), Peroxidase AffiniPure Donkey Anti-Rabbit IgG (Jackson ImmunoResearch, 711-035-152, 1:10,000 dilution).

### NO and nitrite measurement

Intracellular NO levels were measured using OxiSelect Intracellular Nitric Oxide (NO) Assay Kit (Cell Biolabs, STA-800-5). Nitrite levels in the cleared supernatant were measured using Griess Reagent Kit (Thermo Fisher Scientific, G7921) according to the manufacturer’s instructions and detected on a Cytation5 plate reader.

### Plasmids

pLIX-hNos2 (Addgene, 110800), psPAX2 (Addgene, 12260), pMD2.G (Addgene, 12259), LentiGude Puro (Addgene, 52963) were purchased from Addgene. CAS9 Blasticidin Lenti plasmid was purchased from Sigma (CAS9BST). Plasmids were expanded in house and isolated using Plasmid Plus Midi Kit (Qiagen, 12945) according to the manufacturer’s instructions.

sgRNA targeting the respective gene was designed on E-CRISP (http://www.e-crisp.org/) and lentivectors expressing sgRNAs were built according to the published protocol^84^. Primers used to generate the sgRNA plasmids were listed in Supplementary Table S3.

LentiGuide-Gbp-Chr3-sg3+sg4 carrying both Gbp-Chr3 sgRNA3 and sgRNA4 generated by first building individual LentiGuide plasmids as stated above. Amplicon containing U6 promoter and sgRNA4 was PCR amplified using Phusion High-Fidelity DNA Polymerase (New England Biolabs, M0530) and assembled into SalI linearized LentiGuide-GBP-Chr3 sgRNA3 plasmid using HiFi DNA Assembly Kit (New England Biolabs, E5520). Primers used were listed in Supplementary Table S3.

pLIX-Gbp-Chr3-repair plasmid containing a neomycin resistance cassette (*NeoR*) targeting chromosome 3 GBP locus was generated from genomic DNA by amplifying 5’ and 3’ homology arms using primers listed in Supplementary Table S3. The protospacer adjacent motif (PAM) loci of sgRNA were mutated in the amplicons of homology arms. The *NeoR* gene fragment was synthesized by GeneWiz. All fragments were assembled into a pLIX plasmid linearized by NheI and SalI (New England Biolabs) using HiFi DNA Assembly Kit (New England Biolabs, E5520). All plasmids were Sanger sequenced to ensure correct inserts (Eurofins Scientific).

### Lentiviral transduction

HEK-293T cells were seeded 1 day before transfection and transfected with psPAX2, pMD2.G packaging plasmid, and lentivectors using Fugene 6 (Promega, E2691). 24 hours after transfection, media was changed. After 48 hours viral supernatants were collected, cleared of cell debris using 0.45 μm filter (Celltreat, 229766) and added to target cells in the presence of polybrene (EMD Millipore, TR-1003-G). After overnight incubation, fresh media was added. Two days later, cells were selected in 6 μg/mL of puromycin (Sigma, P8833) or 7 μg/mL blasticidin (Research Product International, B122000) for 2 days. Surviving cells were expanded and single-cell cloned.

### Generation of RAW^ΔGbp-chr3^

14 µg of pLIX-Gbp-Chr3-repair plasmid was linearized using NheI and SalI. Purified templates together with 2 µg of LentiGuide-Gbp-Chr3-sg3+sg4 were electroporated into RAW-Cas9 cells using Lonza SF cell line X kit (Lonza, V4XC-2012) on a Lonza 4D-Nucleofector unit (Lonza). Code DS-136 was used for electroporation. Cells were flushed out using warm complete DMEM+10%FBS and rested for 4 days. 1 mg/mL of G418 (Thermo Fisher Scientific, BP6735) was added for 4 days. G418 was added every 2 passages to keep the selection pressure. Successful insertion of the repair template was validated by PCR using primers listed in Supplementary Table S3.

### CellMask staining

Cells and coverslips were prepared as described above. Fixation is followed by a gentle permeabilization with 0.2% (w/v) BSA and 0.02% (w/v) Saponin in PBS (w/o Mg^2+^, Ca^2+^) for 30 minutes at RT and then blocked in PBS+5% BSA for 15 minutes at RT. Primary antibodies were applied for 1h in the permeabilization buffer as described elsewhere. After three 5 minute washes, secondary antibodies were prepared in permeabilization buffer containing 2 μg/ml HCS CellMask Blue Stain (Invitrogen Molecular, H32729) and incubated at RT for 30 minutes protected from light. Following 4 washes, coverslips were mounted with ∼25ul ProLong Glass Antifade Mountant (Thermo, P36980) and left to fully cure overnight in the dark at RT before imaging or longer-term cold storage. The area of the cell mask negative void and the parasite GFP signal was analyzed in Fiji.

### Transmission electron microscopy

5.0x10^6 RAW 264.7 cells were plated into three 10cm dishes and incubated overnight at 37°C with 5% CO2. Subsequently, appropriate dishes were primed with 10ng/ml mIFNγ and Pam3CSK4 for 24 hours prior to infection. An hour before infection, one primed dish was treated with 100 μM 1400W. Both dishes were infected for 6 hours with Type II Me49-GFP-Luc at MOI 2, following which cells were washed thrice with ice-cold PBS.

Simultaneously, a fixation buffer was freshly prepared with 2.5% glutaraledehyde, 0.1M sodium cacodylate (both Sigma-Aldrich, G5882-50ML, and C4945-10G, respectively), and sterile deionized H_2_O. The buffer was applied to the cells, which were then allowed to fix at room temperature for 1 hour. Cells were scraped gently into ’sheets’ and collected into 2-ml Eppendorf tubes, centrifuged, and washed thrice with 0.1M sodium cacodylate. They were stored at 4°C in a 0.1M sodium cacodylate buffer with minimal air exposure.

For ultrastructural analysis, cells were washed with 0.1M sodium cacodylate twice, immersed in 1% osmium tetraoxide (Sigma-Aldrich, 75632) in 0.1M sodium cacodylate at RT for 1 hour, then washed again. Dehydration was carried out using an ethanol gradient (50%, 70%, 95%, and 100%), each for 10 minutes. A series of infiltration steps using an increasing ratio of epoxy resin was executed. Polymerization was performed overnight at 60°C. Ultrathin sections (80 nm) were cut using a Leica EM UC7 Ultramicrotome, collected on 200 mesh copper grids, counterstained with uranyl acetate and lead citrate, and carbon-coated for TEM stability. Samples were stored at RT until imaging. Samples were imaged on a FEI Tecnai F20 equipped with a 4K camera (TSS microscopy). The area of the intravacuolar network (IVN) space was analyzed in Fiji.

### Data availability and analysis

Data were plotted and analyzed in GraphPad Prism 9 and R (version 4.2.0) unless noted otherwise. Raw mass spectrometry data of auoSTOMP (accession: PXD027716) were uploaded to PRIDE (https://www.ebi.ac.uk/pride/).

## Supplemental Information

**Figure S1 (related to Figure 2). *T. gondii* proteins identified by autoSTOMP A.** 86 *T. gondii* proteins were identified using autoSTOMP as described in Figure 2

**Figure S2 (related to Figure 2). Pairwise comparison between autoSTOMP conditions. A**, Pairwise comparison of proteins enriched in the unprimed PVM region (left, negative values) relative to Pam3CSK4 PVM; **B,** IFNγ PVM vs. IFNγ GBP2^+^ vacuoles (middle); **C,** IFNγ+Pam3CSK4 PVM vs. IFNγ+Pam3CSK4 GBP2^+^ vacuoles. Red circles indicate IIG proteins and purple circle indicates iNOS.

**Figure S3 (related to Figure 3). Gating strategy A.** Gating strategy for Figure 3.

**Figure S4 (related to Figure 4). Expression of nitric oxide synthases and IIGs in BM-MoDCs. A**, B6 BM-MoDCs were infected as described in Figure 1. *Nos1* and *Nos3* expression was determined using quantitative reverse transcription polymerase chain reaction (RT-PCR) at 2hpi. N=3 independent experiments. **B,** GBP2 protein levels were detected by Western blot at 14hpi respectively. Volumetric intensity ratios of GBP2 and GAPDH were quantified (right). **C**, IIG expression was determined using quantitative reverse transcription polymerase chain reaction (RT-PCR) at 2 hours post infection. N=3 independent experiments.

**Figure S5 (related to Figure 4). RAW-Cas9 deletion of iNOS and chromosome 3 GBPs. A**, iNOS expression was interrupted by CRISPR/Cas9 in Cas9-expressing RAW 264.7 macrophages and iNOS expression was evaluated in the bulk population or a single cell clone (clone 1) by western blot. **B**, Schematic of the chromosome 3 GBP locus and strategy for generating RAW*^ΔGbp-chr3^* cells using CRISPR-Cas9 mediated homologous recombination. **C** Validation of neomycin resistance cassette integration in chromosome 3 genomic DNA, loss of GBP5 downstream of the integration site and loss GBP2 upstream of the integration site in the bulk population of RAW*^ΔGbp-chr3^* cells (bulk population). **D,** RAW 264.7 macrophages were pretreated for 20 hours with media, 10ng/ml IFNγ and 10ng/ml of Pam3CSK4 for the final 3 hours before infection. Cells were infected with *T. gondii* expressing GFP and luciferase at MOI 5. Cells were pretreated with 1400W 1hr before infection or chloroquine (CHQ) 30 minutes post infection. N=7 independent experiments, 2-way ANOVA with Tuckey post hoc analysis. *, p≤0.05; **, p≤0.01; ****, p≤0.0001.

**Figure S6 (related to Figure 5). iNOS overexpression cannot restrict parasites in the absence of IFNγ. A-B**, RAW 264.7 macrophages were transduced with human *Nos2* under tetracycline inducible promoter and human iNOS expression was titrated by adding no, 200 ng/mL or 400 ng/mL doxycycline which was much lower that OD50 of doxycycline for *T. gondii*. Cells were infected similar to Figure 4 and nitrite production (**A**) was measured to probe for nitric oxide flux while parasite burden was measured by luciferase assay (**B**). N=3 independent experiments, and error bars represent Mean±SEM by 2-way ANOVA with Tukey post hoc analysis. n.s., not significant; *, p≤0.05; **, p≤0.01; ***, p≤0.001; ****, p≤0.0001.

**Figure S7 (related to Figure 5). Inhibiting cellular or mitochondrial ROS does not rescue parasite growth down stream of IFNγ and Pam3CSK4 stimulation, and NO supplementation can not nitrate the vacuole in unstimulated macrophages A-B**, BM-MoDCs were stimulated and infected as described in Figure 1. Parasite growth in mock treated or media supplemented with iNOS inhibitor, 1400W, cellular ROS inhibitor, N-acetyl cysteine (NAC) (**A**) or mitochondrial targeted ROS scavenger, mitoTEMPO (**B**) was measured by luciferase assay at 14 hpi. 2-way ANOVA with Tuckey post hoc analysis. ****, p≤0.0001. **C-D**, RAW 264.7 cells were infected with Me49-GFP-Luc as described in Figure 4. **C,** RNAseq analysis of RAW 264.7 cells at 18hpi. **D,** The NO donor DETA NONOate was added to RAW 264.7 cells one hour prior to infection, and samples were fixed and stained with a nitrotyrosine-specific (NO_2_-Y, magenta) antibody to evaluate co-localization with parasite GFP. Error bars in **C** represent Mean±SEM by 2-way ANOVA with Tukey post hoc analysis. *, p≤0.05; **, p≤0.01; ***, p≤0.001; ****, p≤0.0001.

**Figure S8 (related to Figure 6). PVM ultrastructure of 2 pack vacuole in IFNγ and Pam3CSK4 stimulated RAW 264.7 cells during iNOS inhibition. A**, Vacuole ultrastructure was evaluated by transmission electron microscopy in RAW-Cas9 cells that were unstimulated, treated with IFNγ and Pam3CSK4 with1400W then infected for 6hours. Insets show the intravacuolar network (IVN) between the parasite plasma membrane (*Tg*) and the parasitophorous vacuole membrane (PVM, blue line), * indicates breaks in the PVM, *div. Tg* indicates vacuoles with 2 parasites.

**Table S1. All identified mouse proteins in autoSTOMP**

**Table S2. All identified *T. gondii* proteins in autoSTOMP**

**Table S3. Primers used in this study**

## Acknowledgements

We want to thank the members of Ewald lab for their feedback. We thank Dr. Hervé Agaisse for his insights and suggestions for this manuscript. We thank Dr. André Marette for sharing the *Nos2*^fl/fl^ mice and for constructive discussion of their use to evaluate iNOS biology in vivo. We also thank Dr. Nicholas E. Sherman and the Biomolecular Analysis Facility Core for LC-MS. This work used TEM sample preparation service in the Advanced Microscopy Facility (Research Resource Identifiers (RRID): SCR_018736). Transmission electron micrographs were recorded at the University of Virginia Molecular Electron Microscopy Core facility (RRID:SCR_019031), which is built with NIH grant G20-RR31199. BAF, AMF, and MEMC are supported by the University of Virginia School of Medicine. This work is supported by NIGMS R35GM138381 (S.E.E.) and NIAID R21AI156153 (S.E.E.).

## Notes

### Competing Interest Statement

The authors have declared no competing interest.

