## Supplementary figures and images for "Inducible nitric oxide synthase (iNOS) is necessary for GBP-mediated *T. gondii* restriction in murine macrophages via vacuole nitration and intravacuolar network collapse"

### Supplementary Figures

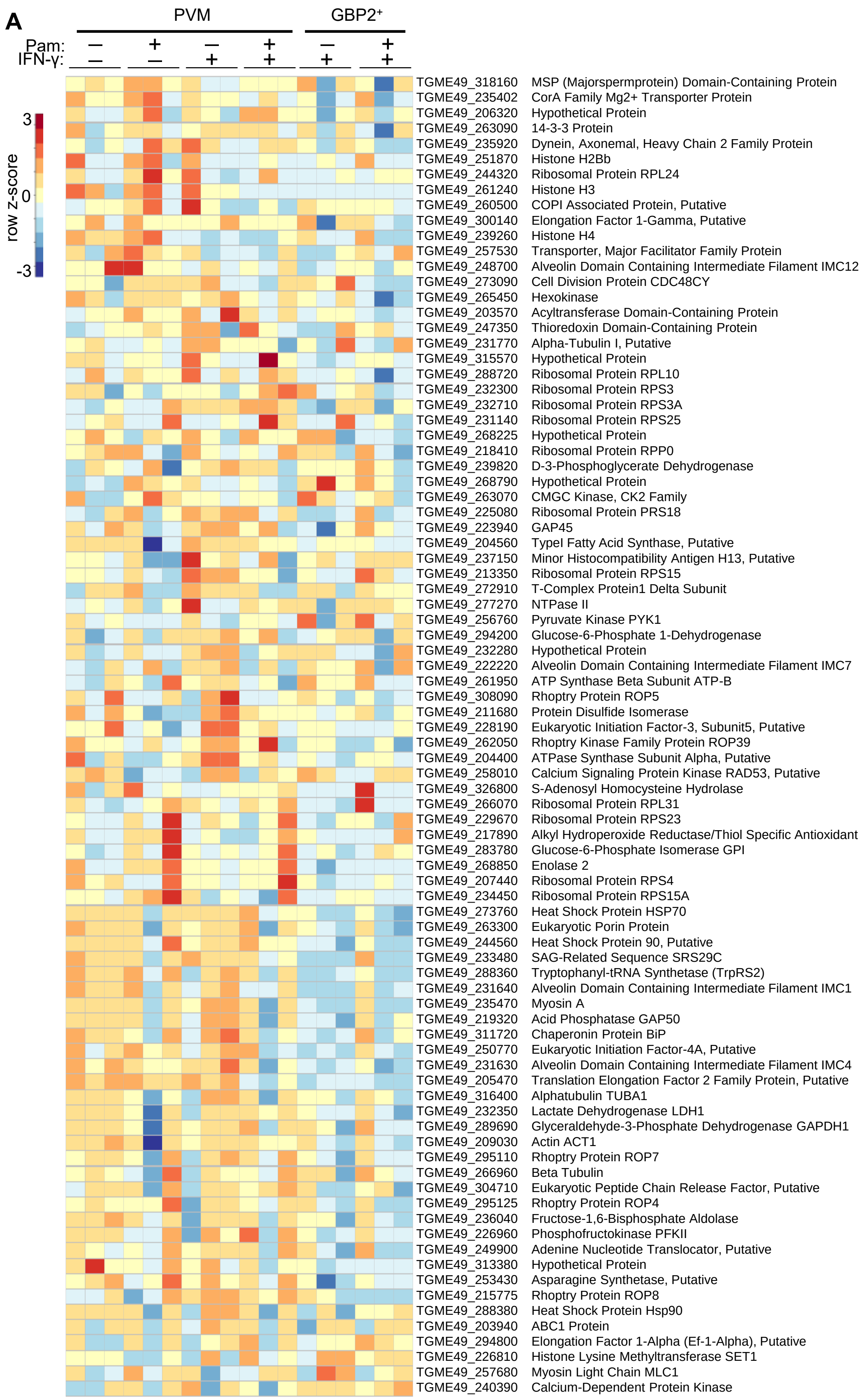

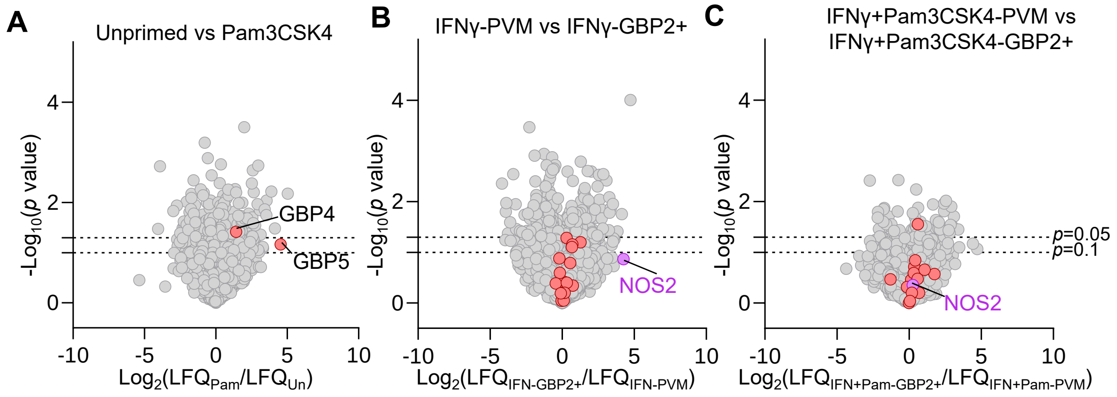

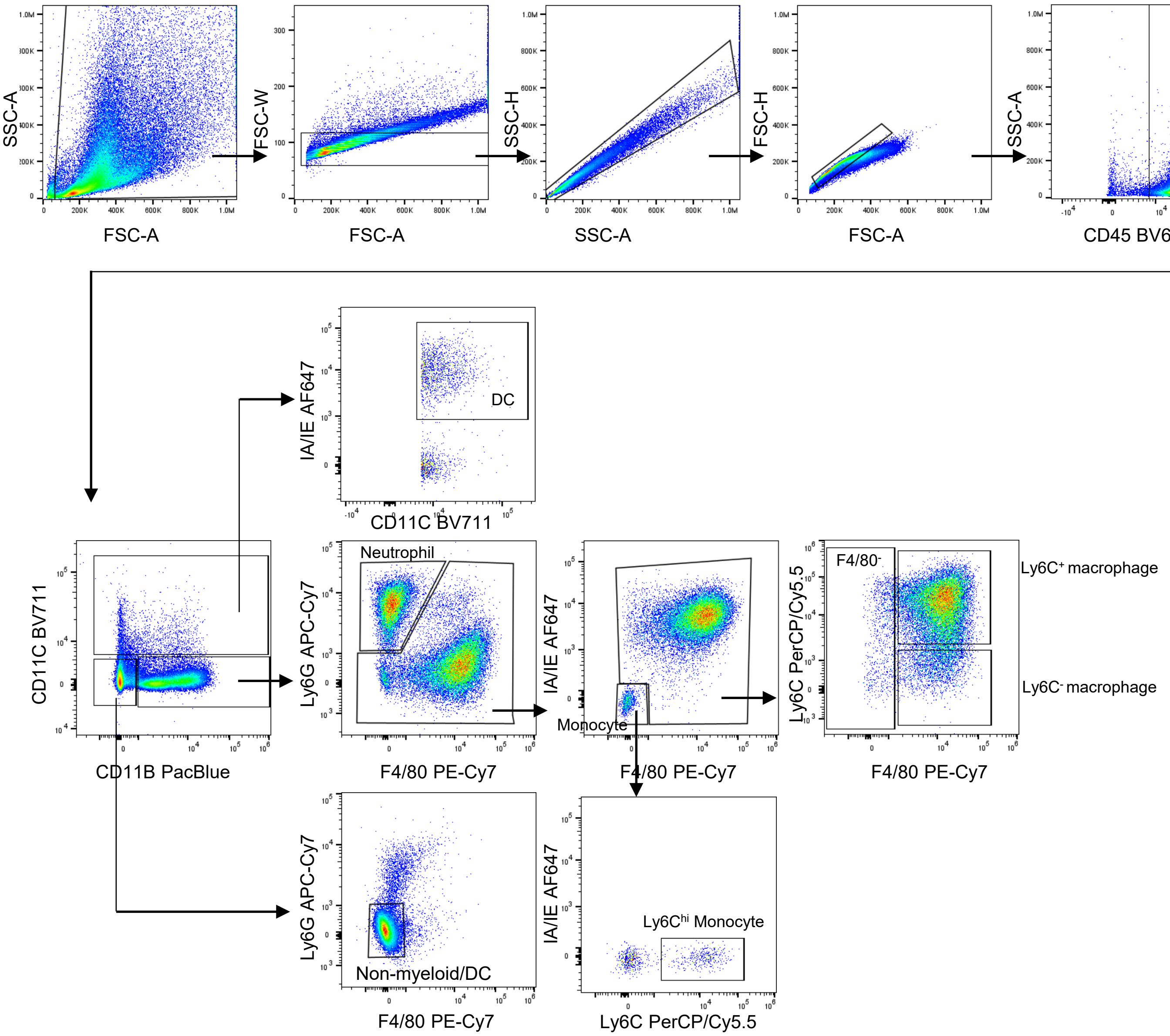

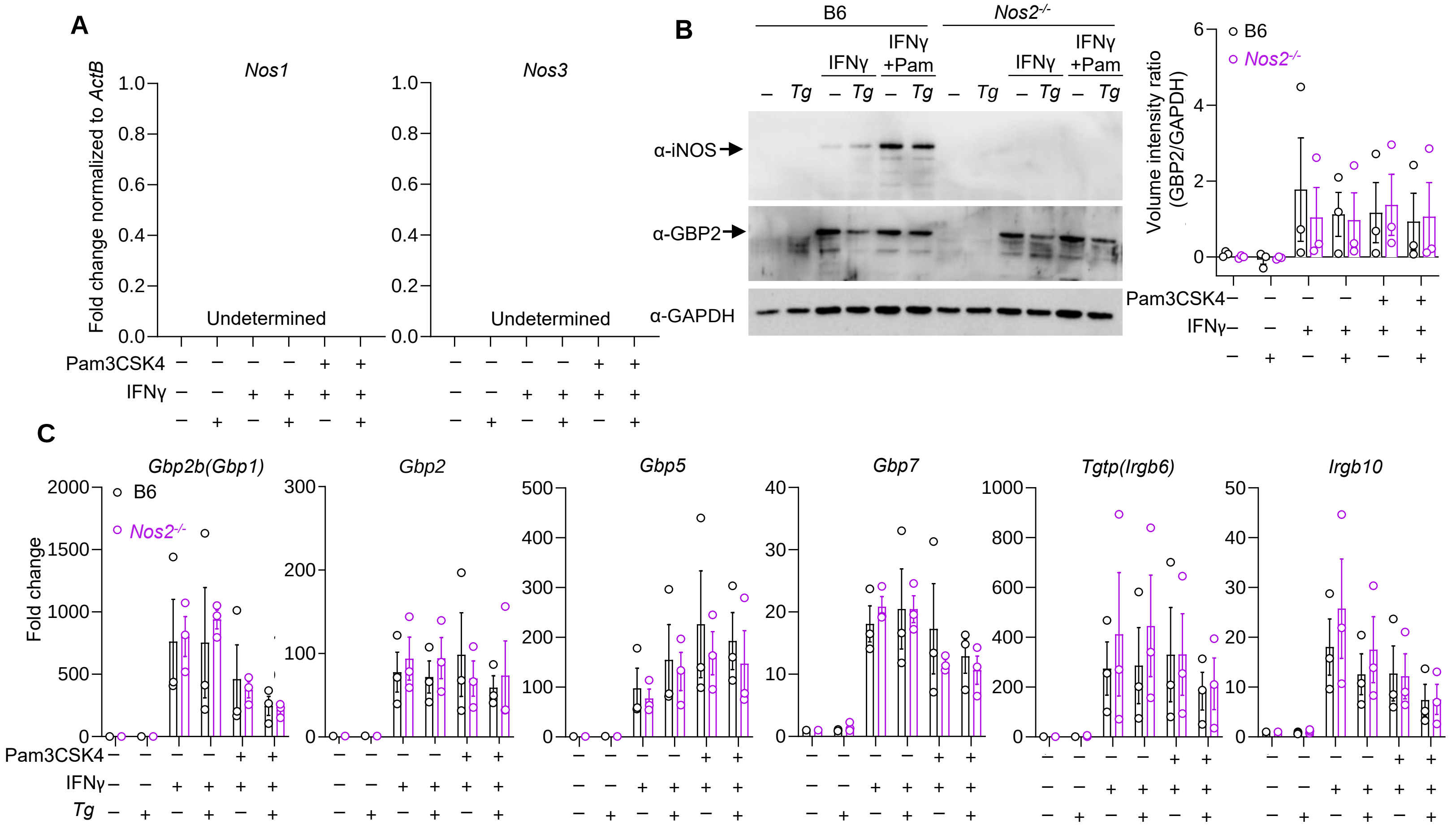

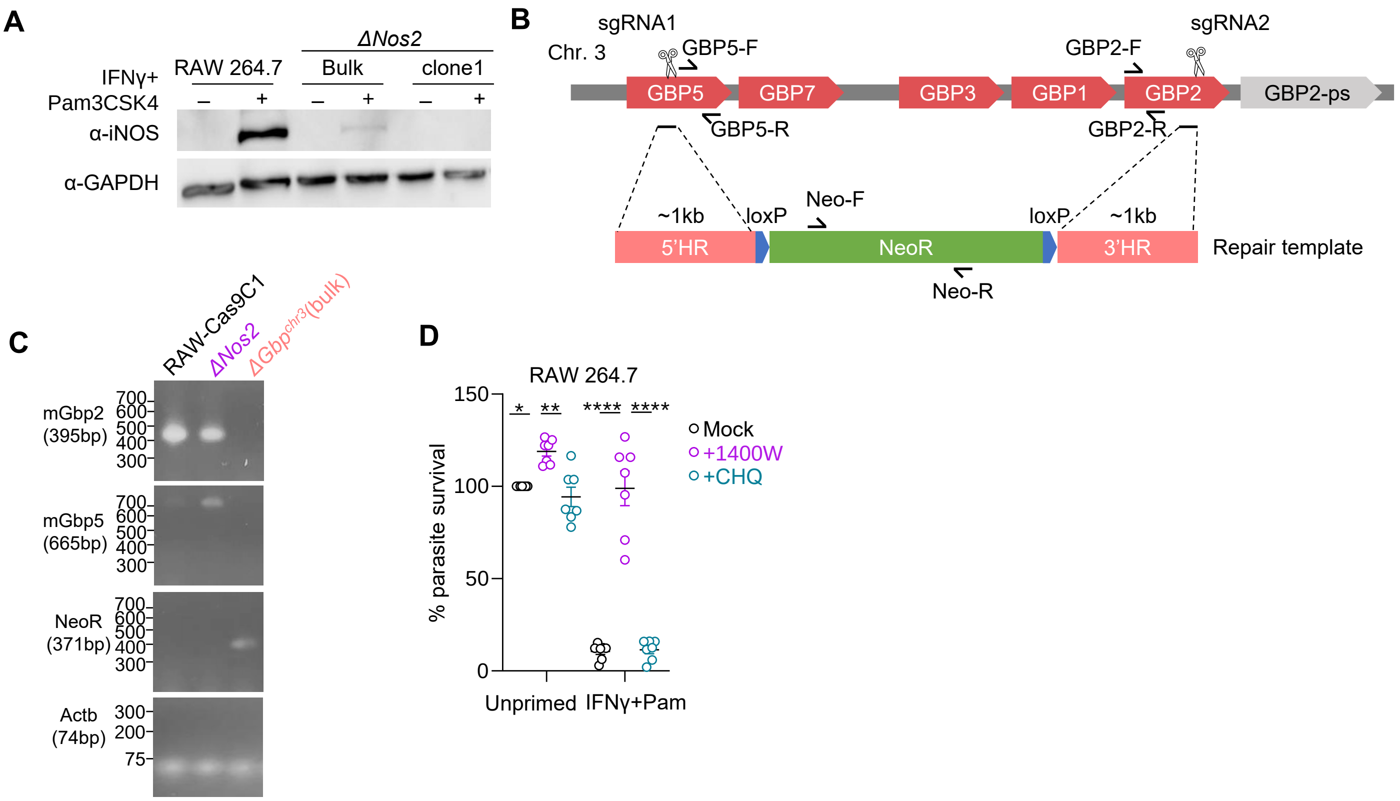

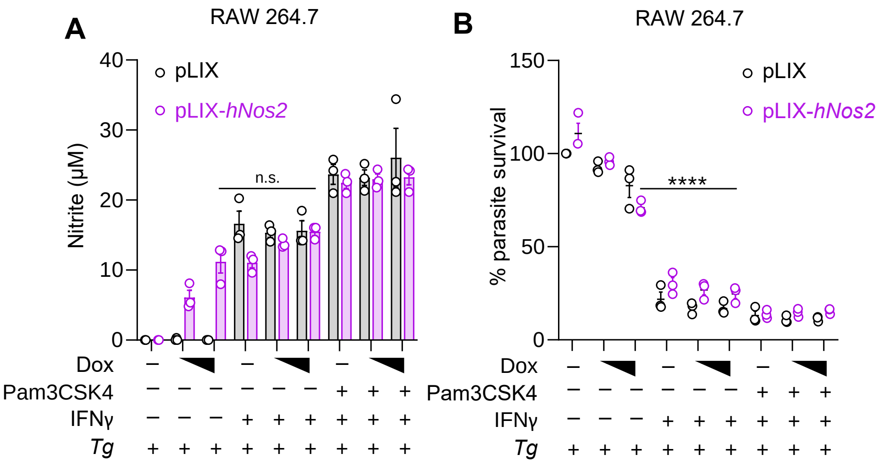

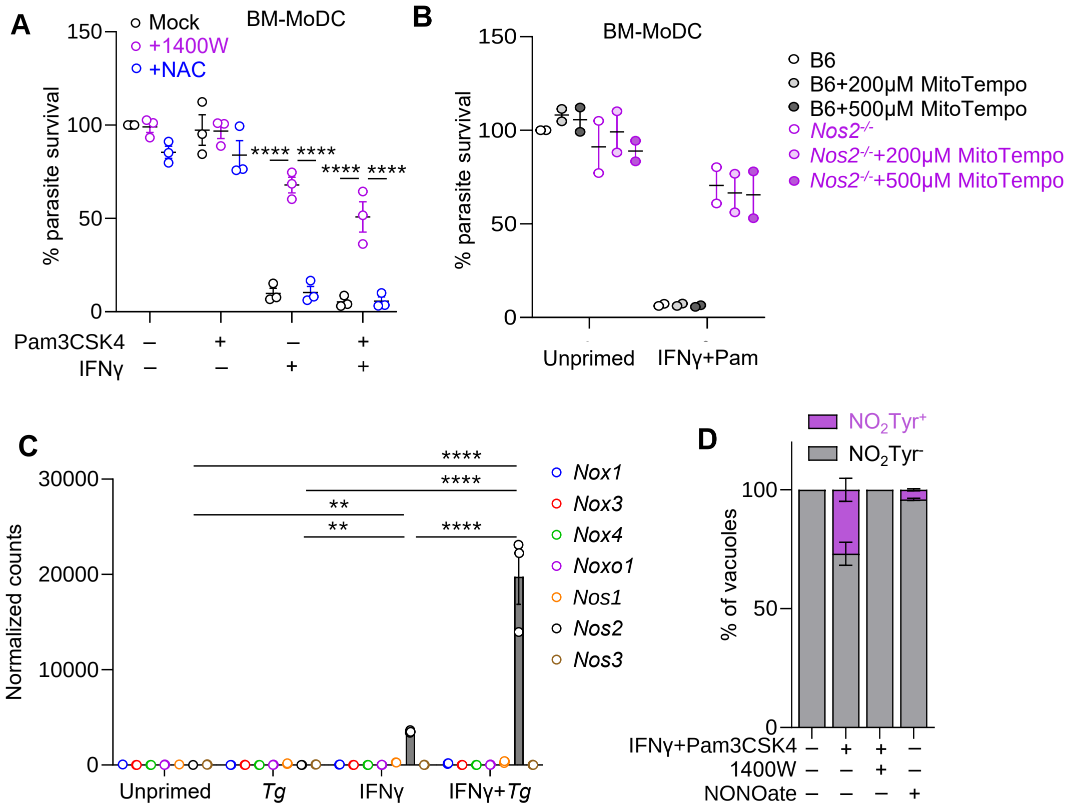

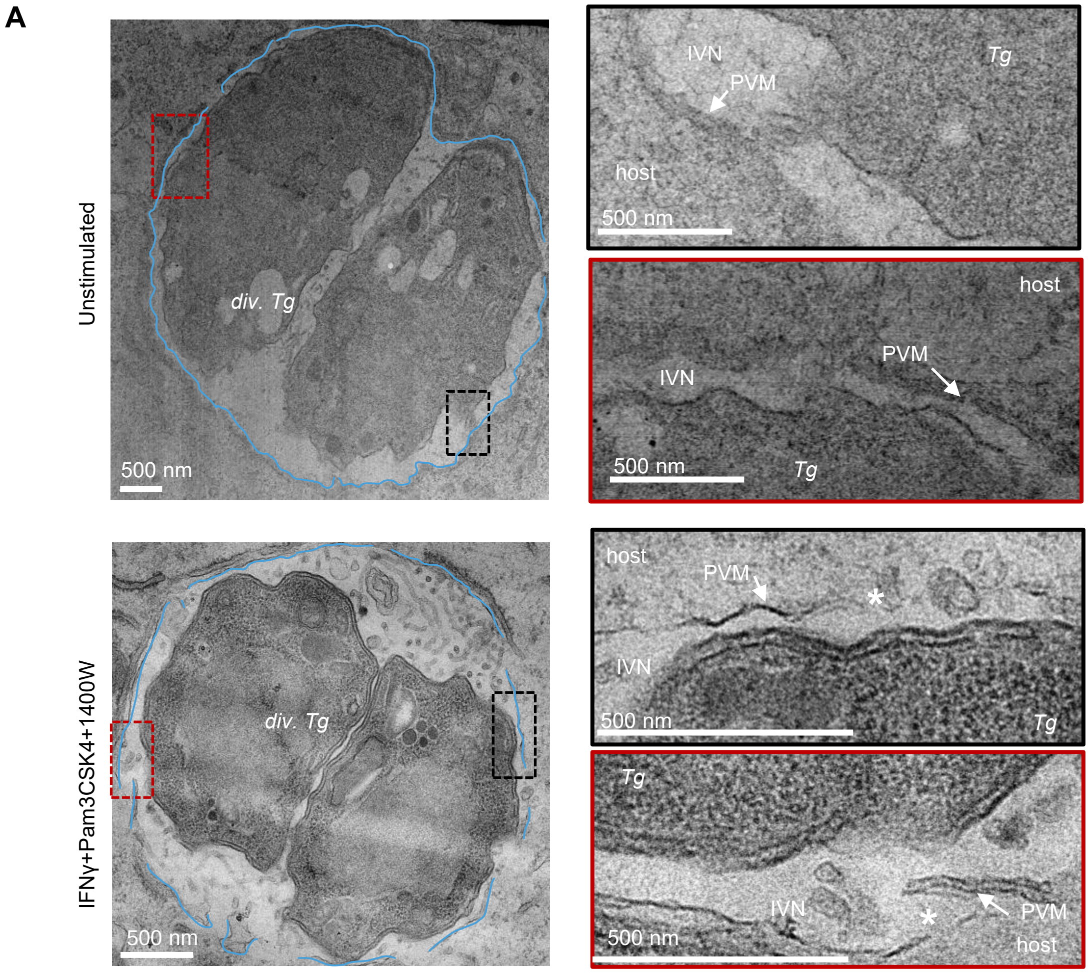
